# The origin of physiological local mGluR1 supralinear Ca^2+^ signals in cerebellar Purkinje neurons

**DOI:** 10.1101/785493

**Authors:** Karima Ait Ouares, Marco Canepari

## Abstract

In Purkinje neurons, the climbing fibre (CF) input provides a signal to parallel fibre (PF) synapses triggering PF synaptic plasticity. This supralinear Ca^2+^ signal, co-localised with the PF Ca^2+^ influx, occurs when PF activity precedes the CF input. Using membrane potential (V_m_) and Ca^2+^ imaging, we identified the biophysical determinants of these supralinear Ca^2+^ signals. The CF-associated Ca^2+^ influx is mediated by T-type or by P/Q-type Ca^2+^ channels, depending on whether the dendritic V_m_ is hyperpolarised or depolarised. The resulting Ca^2+^ elevation is locally amplified by saturation of the endogenous Ca^2+^ buffer produced by the PF-associated Ca^2+^ influx, in particular by the slow Ca^2+^ influx mediated by type-1 metabotropic glutamate receptors (mGluR1s). When the dendrite is hyperpolarised, mGluR1s boost neighbouring T-type channels providing a mechanism for local coincident detection of PF-CF activity. In contrast, when the dendrite is depolarised, mGluR1s increase dendritic excitability by inactivating A-type K^+^ channels, but this phenomenon is not restricted to the activated PF synapses. Thus, V_m_ is likely a crucial parameter in determining PF synaptic plasticity and the occurrence of hyperpolarisation episodes is expected to play an important role in motor learning.

## INTRODUCTION

In cerebellar Purkinje neurons (PNs), short- and long-term synaptic plasticity of the parallel fibre (PF) inputs can be induced by pairing PF excitatory synaptic potentials (EPSPs) with a climbing fibre (CF) EPSP (Wang et al., 2000; Brenowitz and Regehr, 2005; Safo and Regehr, 2005). These forms of plasticity, that are believed to underlie associative motor learning and control behaviours (Ito, 2001), are homosynaptic and they require local Ca^2+^ elevation at the site of activated PF synapses. Precisely, the Ca^2+^ transient triggered by a CF-EPSP alone is not local (Canepari and Vogt, 2008; Ait Ouares et al., 2019) since it is caused by a depolarising transient, originating in the soma and in the initial dendritic segment, that spreads to distal dendrites to open voltage-gated Ca^2+^ channels (VGCCs). When a CF-EPSP occurs after PF activity, the associated Ca^2+^ transient is larger than the summation of the two Ca^2+^ transients triggered by PF and CF inputs alone, and it is therefore referred as “supralinear” Ca^2+^ signal (Brenowitz and Regehr, 2005). However, in contrast to the Ca^2+^ transient associated with a CF-EPSP alone, the supralinear Ca^2+^ signal associated with paired PF and CF inputs is localised exclusively at the site of activated PF synapses.

The ability of a spread physiological signal to trigger a local signal, in combination with synaptic activity, exclusively at the sites of activated synapse, is a crucial requirement for coincident detection and associative plasticity in many systems. In hippocampal and neocortical pyramidal neurons, synchronous action potentials and excitatory synaptic activity can potentiate activated synapses (Debanne, 2001). In this case, action potentials transiently depolarising the dendrites can provide a signal localised at activated synapses by unblocking NMDA receptors from Mg^2+^ (Ascher and Nowak, 1987). In the case of PF synaptic plasticity induced by coincident CF inputs, however, the underlying biophysical mechanism does not involve NMDA receptors but instead type-1 metabotropic glutamate receptors (mGluR1s) which are locally activated by glutamate release from PF terminals (Hartell, 1994; Wang et al., 2000; Brenowitz and Regehr, 2005; Safo and Regehr, 2005). Since mGluR1 activation is not voltage-dependent, the fundamental question is to understand how a CF-EPSP, that alone generates a depolarisation and a Ca^2+^ influx everywhere in the dendrite, can trigger a strong localised Ca^2+^ elevation, in combination with mGluR1 activation, exclusively at the site of activated PFs.

In a recent study, we have demonstrated that the dendritic depolarisation produced by a CF-EPSP activates two types of VGCCs, namely T-type and P/Q-type VGCCs, and that the activation of both channels is determined by the dendritic membrane potential (V_m_) at the beginning of the CF-EPSP (Ait Ouares et al., 2019). Specifically, at initial hyperpolarised V_m_, the dendritic Ca^2+^ transient is largely mediated by T-type VGCCs while activation of A-type VGCCs limits the opening of P/Q-type VGCCs. In contrast, at initial depolarised V_m_, T and A channels are inactivated and the larger dendritic depolarising transient activates P/Q-type VGCCs that mediate Ca^2+^ spikes. It was also shown that both T-type VGCCs (Hildebrand et al., 2009; Isope et al., 2012) and P/Q-type VGCCs (Otsu et al., 2014) can be boosted by mGluR1 activation, providing two potential biophysical mechanisms underlying localised supralinear Ca^2+^ signals that are required for coincident detection and PF synaptic plasticity.

In the present study we addressed the question of how a CF-EPSP can generate a supralinear Ca^2+^ signal exclusively at the sites of activated PF synapses. We used a protocol of moderate PF stimulation mimicking a physiological burst to reveal the biophysical mechanism that permit a CF-EPSP to trigger a localised signal at the site of activated PF synapses in realistic scenarios. We used ultrafast V_m_ and Ca^2+^ imaging, using indicators of different affinity and a series of selective pharmacological manipulations and we performed a detailed analysis to elucidate the different mechanisms underlying supralinear Ca^2+^ signals under different conditions.

## MATERIALS AND METHODS

### Slice preparation, solutions, electrophysiology and pharmacology

Experiments were ethically carried out in accordance with European Directives 2010/63/UE on the care, welfare and treatment of animals. Procedures were reviewed by the ethics committee affiliated to the animal facility of the university (D3842110001). We used 21-35 postnatal days old mice (C57BL/6j). Animals were anesthetised by isofluorane inhalation and the entire cerebellum was removed after decapitation. Cerebellar sagittal slices (250 µm thick) were prepared following established procedures (Vogt et al., 2011a; Vogt et al., 2011b; Ait Ouares et al., 2016) using a Leica VT1200 (Leica, Wetzlar, Germany). Slices were incubated at 37°C for 45 minutes before use. The extracellular solution contained (in mM): 125 NaCl, 26 NaHCO_3_, 1 MgSO_4_, 3 KCl, 1 NaH_2_PO_4_, 2 CaCl_2_ and 20 glucose, bubbled with 95% O_2_ and 5% CO_2_. The intracellular solution contained (in mM): 125 KMeSO_4_, 5 KCl, 8 MgSO_4_, 5 Na_2_-ATP, 0.3 Tris-GTP, 12 Tris-Phosphocreatine, 20 HEPES, adjusted to pH 7.35 with KOH. In V_m_ imaging experiments, PNs were loaded with the voltage-sensitive dye (VSD) JPW1114 as previously described (Canepari and Vogt, 2008) and in some experiments also with the Ca^2+^ indicator Fura-2FF (at 1 mM) using a previously described procedure (Vogt et al., 2011a). In experiments of Ca^2+^ imaging only, Oregon Green BAPTA-5N (OG5N) was added to the internal solution at 2 mM concentration and, in experiments reported in Figure 5, also Fura-2 (Fura2) at 0.4 mM concentration was included. Patch-clamp recordings were made at 32-34°C using a Multiclamp amplifier 700A (Molecular Devices, Sunnyvale, CA) and electrical signals were acquired at 20 kHz using the A/D board of the CCD camera, regardless of the imaging acquisition rate. The measured V_m_ was corrected for junction potential (−11 mV) as previously estimated (Canepari et al., 2010). PF- and CF-EPSPs were elicited by current pulses of 5-20 µA amplitude and 100 µs duration delivered by separate electrodes. All recordings were performed at least 30 minutes after establishing whole. This minimum time before starting recording was 45 minutes in experiments with Fura2, heparin and ryanodine. In V_m_ imaging experiments, cells were initially loaded with the voltage sensitive dye JPW1114 and re-patched with a solution without the dye to avoid toxicity due to dye overloading. 7-(Hydroxyimino)cyclopropa[*b*]chromen-1a-carboxylate ethyl ester (CPCCOEt), (1*S*,2*S*)-2-[2-[[3-(1*H*-Benzimidazol-2-yl)propyl]methylamino]ethyl]-6-fluoro-1,2,3,4-tetrahydro-1-(1-methylethyl)-2-naphthalenyl-cyclopropanecarboxylate-dihydrochloride (NNC550396 or NNC) and *N*,*N*,*N*-Trimethyl-5-[(tricyclo[3.3.1.1^3,7^]dec-1-ylmethyl)amino]-1-pentanaminium bromide hydrobromide (IEM1460) were dissolved in exracellular solution and applied to the bath *via* the perfusion system. Heparin sodium salt and 1*H*-Pyrrole-2-carboxylic acid, (3*S*,4*R*,4a*R*,6*S*,7*S*,8*R*,8a*S*,8b*R*,9*S*,9a*S*)-dodecahydro-4,6,7,8a,8b,9a-hexahydroxy-3,6a,9-trimethyl-7-(1-methylethyl)-6,9-methanobenzo[1,2]pentaleno[1,6-*bc*]furan-8-yl ester (ryanodine) were dissolved in intracellular solution and applied *via* the patch recording. DHPG, dissolved in external solution, was applied through a pipette positioned near the PN dendrite using 20 ms pulses of pressure provided by a pressure ejector PDES-2DX (npi, Tamm, Germany). AmmTx, dissolved in external solution, was applied through a pipette positioned near the PN dendrite by continuous gentle pressure application. CPCCOEt, Heparin sodium salt, ryanodine, NNC550396 and DHPG were purchased by Tocris (Bristol, UK). IEM1460 was purchased by Hello Bio (Bristol, UK). AmmTx was purchased by Smartox (Saint Egrève, France).

### Optical recording

OG5N (Invitrogen, Carlsbad, CA) was excited at 470 nm with an OptoLED (Jaafari et al., 2014; Jaafari and Canepari, 2016; Ait Ouares et al., 2016; Ait Ouares et al., 2019). Fura-2FF (Santa Cruz, Dallas, TX) and Fura-2 (Invitrogen) were excited at 385 nm using the OptoLED (Cairn Research, Faversham, UK). V_m_ and Ca^2+^ optical measurements were achieved sequentially (Canepari and Ogden, 2008) by alternating excitation of Fura-2FF excitation and the excitation of the voltage sensitive dye JPW1114 (Invitrogen) at 528 nm using a LDI laser (89 North, Williston, VT). OG5N Ca^2+^ fluorescence was recorded at 530 ± 21 nm. Fura-2FF and Fura2 Ca^2+^ fluorescence was recorded at 510 ± 41 nm. JPW1114 V_m_ fluorescence was recorded at >610 nm. Image sequences were recorded using a NeuroCCD-SMQ camera (RedShirtimaging, Decatur, GA). Images were de-magnified by ∼0.2X to visualise an area of ∼150 µm diameter and acquired at 5 kHz or at 20 kHz with a resolution of 26 × 26 pixels or 26 × 4 pixels respectively. In all experiments, four trials with 20 s between two consecutive trials, were obtained to assess the consistency of the signals and averaged to improve the signal-to-noise ratio. The number of averaged trials was nine in 20 kHz recordings. However, in all figures, the somatic V_m_ recording used in the ilustrations was from a single trial, corresponding to the first trial of the series. Fluorescence was corrected for bleaching using a filtered trial without signal. Ca^2+^ fluorescence transients were expressed as fractional changes of fluorescence (ΔF/F_0_).

### Experimental design and statistical analysis

Data were processed and analysed using MATLAB. Ca^2+^ fluorescence transients were expressed as fractional changes of fluorescence (ΔF/F_0_). V_m_ fluorescence transients were calibrated in mV using prolonged hyperpolarising step as described in a previous article (Ait Ouares et al., 2019). Changes produced by the pairing protocol with respect to the CF stimulation alone, or by the pharmacological action of an agent, were assessed by performing the paired Student’s t-test on the signal parameter under the two different conditions. A change in the signal was considered significant when p < 0.01. In all figures, a significant change was indicated with “*”. In the specific case of data reported in Figure 1, the conclusion that the effect of adding CPCCOEt is different when the CF stimulation is delayed by 60 ms from the first PF stimulus, with respect to the cases of 100 ms and 150 ms delays, is supported by a paired t-test performed on the ΔF/F_0_ ratios at two different delays that gave a value of p < 0.01.

**Figure 1.**
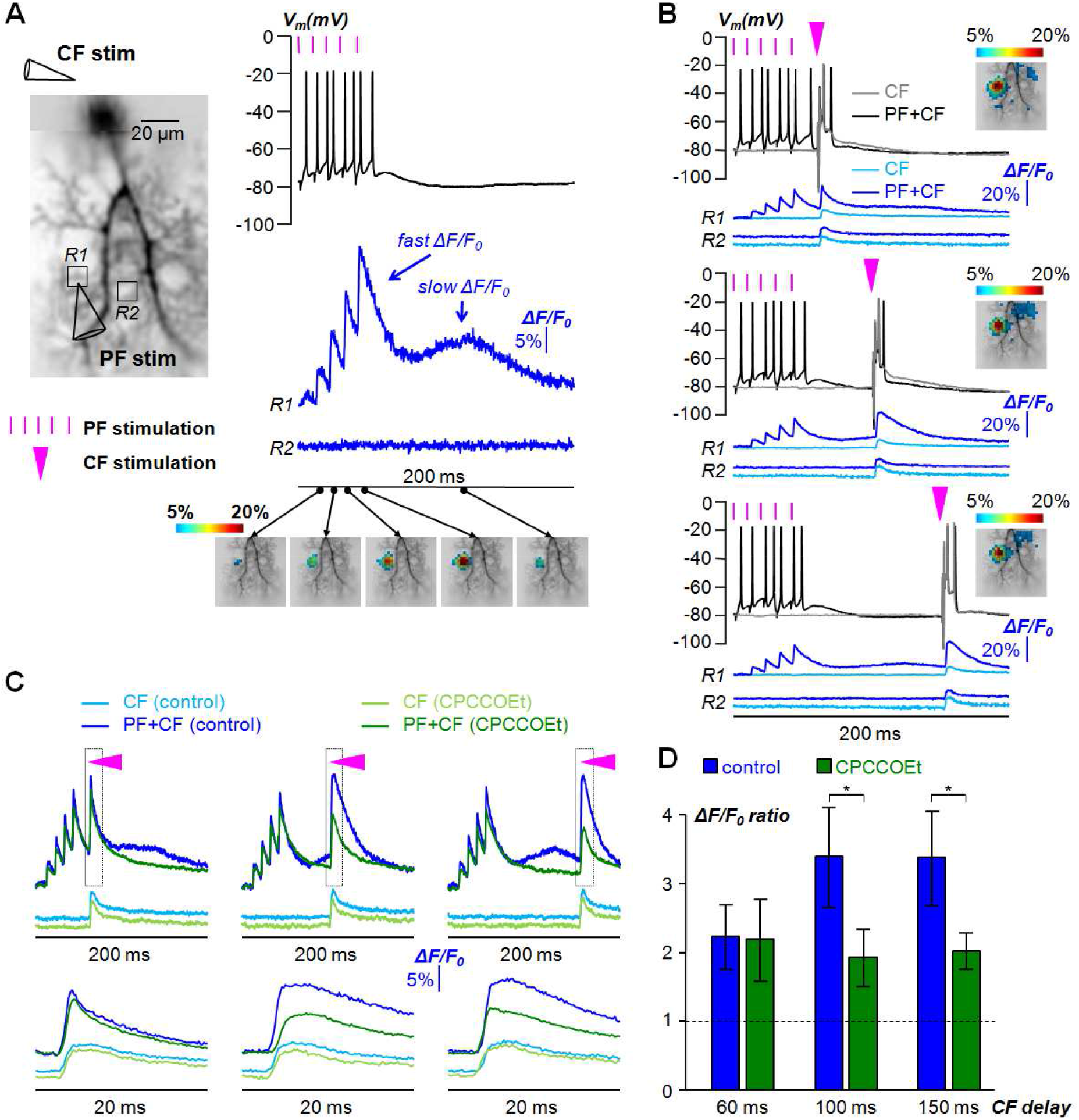
Timing and localisation of supralinear Ca^2+^ signals. (A) Reconstruction of PN filled with 2 mM OG5N (left) with position of two stimulating electrodes for the CF and PFs and two regions of interest indicated (*R1* and *R2*). Somatic V_m_ and Ca^2+^ ΔF/F_0_ signals at *R1* and *R2* associated PFs stimulated with 5 pulses at 100 Hz with timing indicated by purple lines (right). Spatial distribution of ΔF/F_0_ signal corresponding to the 2^nd^-5^th^ PF-EPSPs and to the peak of the slow Ca^2+^ signal depicted using a colour code. (B) Somatic V_m_ and Ca^2+^ ΔF/F_0_ signals at *R1* and *R2* associated with CF stimulation alone (light blue traces) or paired (dark blue traces) with PF stimulation at 100 Hz (timing indicated by purple lines) following at 60 ms, 100 ms or 150 ms from the first PF pulse (timing indicated by purple triangle). Spatial distribution of the ΔF/F_0_ signal associated with CF-EPSP paired with PF stimulation depicted using a colour code. (C) Ca^2+^ ΔF/F_0_ signals at *R1* associated with CF stimulation alone or paired with PF stimulation delayed by 60 ms, 100 ms or 150 ms from the first PF pulse in control solution (blue traces) or after addition of 20 µM of the mGluR1 antagonist CPCCOEt (green traces). The time window outlined is reported below. (D) Mean ± SD (N = 6 cells) of ΔF/F_0_ ratio between the paired and unpaired signals in control solution or after addition of CPCCOEt. “*” indicates significant inhibition of supralinear Ca^2+^ signal by CPCCOEt (p< 0.01, paired t-test). The dotted line depicts ΔF/F_0_ ratio = 1.

For the quantitative interpretation of the results of combined Ca^2+^ imaging with the low-affinity indicator OG5N and with the high affinity indicator Fura2, we calculated the variable “*S*” defined as:

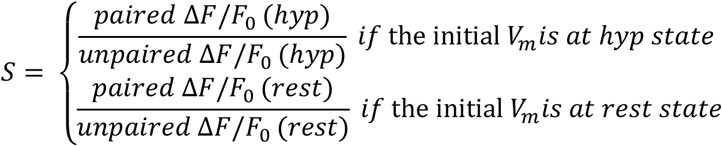

It must be noticed that *S* is principally used to analyse the supralinear Ca^2+^ signal at *hyp* and at *rest* state, and therefore both the numerator the denominator are different according to the different initial V_m_. If a supralinear Ca^2+^ signal occurs when a CF-EPSP is evoked under two different conditions, for instance unpaired and paired, then *S*(OG5N) is obviously positive since a larger free Ca^2+^ concentration implies a larger fraction of Ca^2+^ bound to the low-affinity indicator OG5N. The parameter *S*(Fura2) can unambiguously discriminate whether a supralinear Ca^2+^ signal is or is not exclusively due to the saturation of Fura2, and whether it is or it is not exclusively due to a larger Ca^2+^ influx through the plasma membrane.

1. *Case of supralinear Ca^2+^ signal exclusively due to the saturation of Fura2*. In the presence of Fura2, the larger fraction under paired conditions can be in principle exclusively due to the saturation of the high-affinity indicator by the Ca^2+^ associated with the PF stimulation. Thus, if less free Fura2 is available to buffer the Ca^2+^ entering the cell during the CF-EPSP, more Ca^2+^ will bind to OG5N and less Ca^2+^ will bind to Fura2 under paired conditions, implying that *S*(Fura2) will be smaller than 1. Thus, if *S*(Fura2) is larger than 1, it unambiguously proves that the supralinear Ca^2+^ signals is not exclusively due to the saturation of Fura2, i.e. that at least a fraction Ca^2+^ bound to the low-affinity indicator OG5N originates from Ca^2+^ influx through the plasma membrane.
2. *Case of supralinear Ca^2+^ signal exclusively due to Ca^2+^ influx through the plasma membrane*. If the concentration of Fura2 bound to Ca^2+^ associated with the PF stimulation is negligible with respect to the free Fura2 concentration at the time of the CF-EPSP, then the Ca^2+^ transient entering the cell will linearly bind to OG5N and to Fura2, implying that *S* will have the same value for the two indicators. Thus, if the ratio *S*(Fura2)/*S*(OG5N) = 1, this means that Fura2 is not saturated. Alternatively, if *S*(Fura2)/*S*(OG5N) < 1, this means that at least a fraction Ca^2+^ bound to the low-affinity indicator OG5N originates from the saturation of Fura2.

In the specific case in which the unpaired Ca^2+^ signal is the same, i.e. when the initial V_m_ is at the *hyp* state and the delay of the CF stimulation is either 60 ms or 110 from the first PF stimulus, the ratio *S*(Fura2)/*S*(OG5N) provides a comparative estimate of the Fura2 saturation. Indeed, regardless of the contribution of the Ca^2+^ influx that is different at the two different delays, the more Fura2 is saturated the smaller *S*(Fura2) will be and, conversely, the larger *S*(OG5N) will be. Thus, the smaller *S*(Fura2)/*S*(OG5N) is, the larger the contribution of Fura2 saturation will be.

## RESULTS

### Distint dendritic supralinear Ca^2+^ transients associated with paired PF-CF stimulation

Several studies have shown that, at sites of activated PF synapses, the Ca^2+^ transient associated with a CF-EPSP is significantly larger when it occurs after PF activation (Wang et al., 2000; Brenowitz and Regehr, 2005; Safo and Regehr, 2005; Canepari and Vogt, 2008). To initially analyse these supralinear Ca^2+^ signals, we used a protocol of stimulation for the PF input consisting of 5 stimuli delivered at 100 Hz with an electrode positioned near a PN dendritic site (Figure 1A), with the initial V_m_ hold between −75 mV and −85 mV, below the average V_m_ recorded in the soma when no current is injected (V_rest_), which was always between −65 mV and −50 mV. The intensity of stimulation was set to attain a first EPSP in the range of 1-4 mV of somatic amplitude and the protocol occasionally caused somatic action potentials during the last EPSPs. In cells filled with 2 mM of the low-affinity Ca^2+^ indicator OG5N (K_D_ = 35 µM, Canepari and Ogden, 2006), this protocol systematically produced a fast fractional change of fluorescence (ΔF/F_0_) signal, raising at each individual PF-EPSP, followed by a slow ΔF/F_0_ signal peaking at 100-150 ms after the first PF stimulus. As illustrated by the colour code pictures in Figure 1A, both signals were localised within the same area of a few microns adjacent to the PF stimulating electrode. At submicron spatial scale, however, the two signals are probably not co-localising, since the fast signal is mediated by VGCCs activated by the PF depolarisation (Canepari and Vogt, 2008), whereas the slow signal is mediated by a non selective cation conductance (Canepari et al., 2004) which is believed to be formed by TRPC3 channels (Hartmann et al., 2008).

The time course of the Ca^2+^ transient in the 2X2 pixels (∼11X11 µm^2^) region (*R1*) adjacent to the PF stimulating electrode is reported together with ΔF/F_0_ from a control region (*R2*) of the same size with no Ca^2+^ signal to show the high signal-to-noise ratio of these measurements permitting a precise quantitative analysis of Ca^2+^ signals. We investigated the Ca^2+^ signal associated with a CF-EPSP alone or paired with the PF-EPSPs at delays of 60 ms, 100 ms or 150 ms from the first PF stimulus (Figure 1B). In all three cases, the Ca^2+^ transient associated with the paired CF-EPSP was larger than that associated with the CF-EPSP alone and these phenomena were restricted to the area where the two Ca^2+^ transients associated with the PF-EPSPs were observed, as illustrated by the colour code frames in Figure 1B. Addition of the mGluR1 inhibitor CPCCOEt (20 µM) reduced the slow ΔF/F_0_ signal associated with PF stimulation (Figure 1C) and the supralinear Ca^2+^ signals associated with the CF-EPSP delayed by 100 ms and 150 ms from the first PF stimulus, but not that associated with the CF-EPSP delayed by 60 ms from the first PF stimulus. In N = 6 cells, we quantified the supralinear Ca^2+^ signal by calculating the ratio between the amplitude of the ΔF/F_0_ signal associated with the paired CF-EPSP and the amplitude of the ΔF/F_0_ signal associated with the CF-EPSP alone (ΔF/F_0_ ratio). The ΔF/F_0_ ratio significantly (p < 0.01, paired t-test) decreased from 3.40 ± 0.73 to 1.93 ± 0.42 and from 3.38 ± 0.69 to 2.02 ± 0.27 in the cases of 100 ms and 150 ms delay of the CF stimulation respectively. In contrast, it did not significantly (p > 0.1, paired t-test) change (from 2.23 ± 0.47 to 2.19 ± 0.60) in the case of 60 ms delay of the CF stimulation. We concluded that supralinear Ca^2+^ signals associated with CF-EPSPs delayed by 100 ms and 150 ms were mGluR1-dependent, whereas the supralinear Ca^2+^ signal associated with the CF-EPSP delayed by 60 ms was mGluR1-independent.

The CF-associated Ca^2+^ transient at initial hyperpolarised V_m_ (*hyp*, ∼-80 mV) is mainly mediated by Ca^2+^ influx *via* T-type VGCCs (Ait Ouares et al., 2019). Thus, we further investigated the supralinear Ca^2+^ signal at V_rest_ (*rest*, ∼-60 mV), i.e. when T-type VGCCs are mostly inactivated and the CF-associated Ca^2+^ transient is largely mediated by Ca^2+^ influx *via* P/Q-type VGCCs that can be activated because also most of A-type K^+^ channels are inactivated. Again, we used the protocol consisting of 5 PF stimuli at 100 Hz with an electrode positioned near a PN dendritic site (Figure 2A), followed by a CF-EPSP delayed by 110 ms from the first PF stimulus. In this experiment, however, recordings were performed both at *hyp* and *rest* initial V_m_ (Figure 2B). In both cases, the Ca^2+^ transient associated with the paired CF-EPSP was larger than that associated with the CF-EPSP alone, and both supralinear Ca^2+^ signals were co-localised with the PF active area (see the colour code frames in Figure 2B). Addition of CPCCOEt reduced both supralinear Ca^2+^ signals associated with the CF-EPSP (Figure 2C). In N = 6 cells, the ΔF/F_0_ ratio significantly (p < 0.01, paired t-test) decreased from 3.02 ± 0.61 to 1.88 ± 0.50 and from 2.19 ± 0.38 to 1.46 ± 0.30 in the cases of *hyp* and *rest* initial V_m_ respectively (Figure 2D). We concluded that supralinear Ca^2+^ signals associated with CF-EPSPs, delayed by 110 ms from the first PF stimulus, were mGluR1-dependent both at *hyp* and at *rest* initial V_m_, i.e. independently on whether the CF-associated Ca^2+^ transient is mediated by T-type or P/Q-type VGCCs.

**Figure 2.**
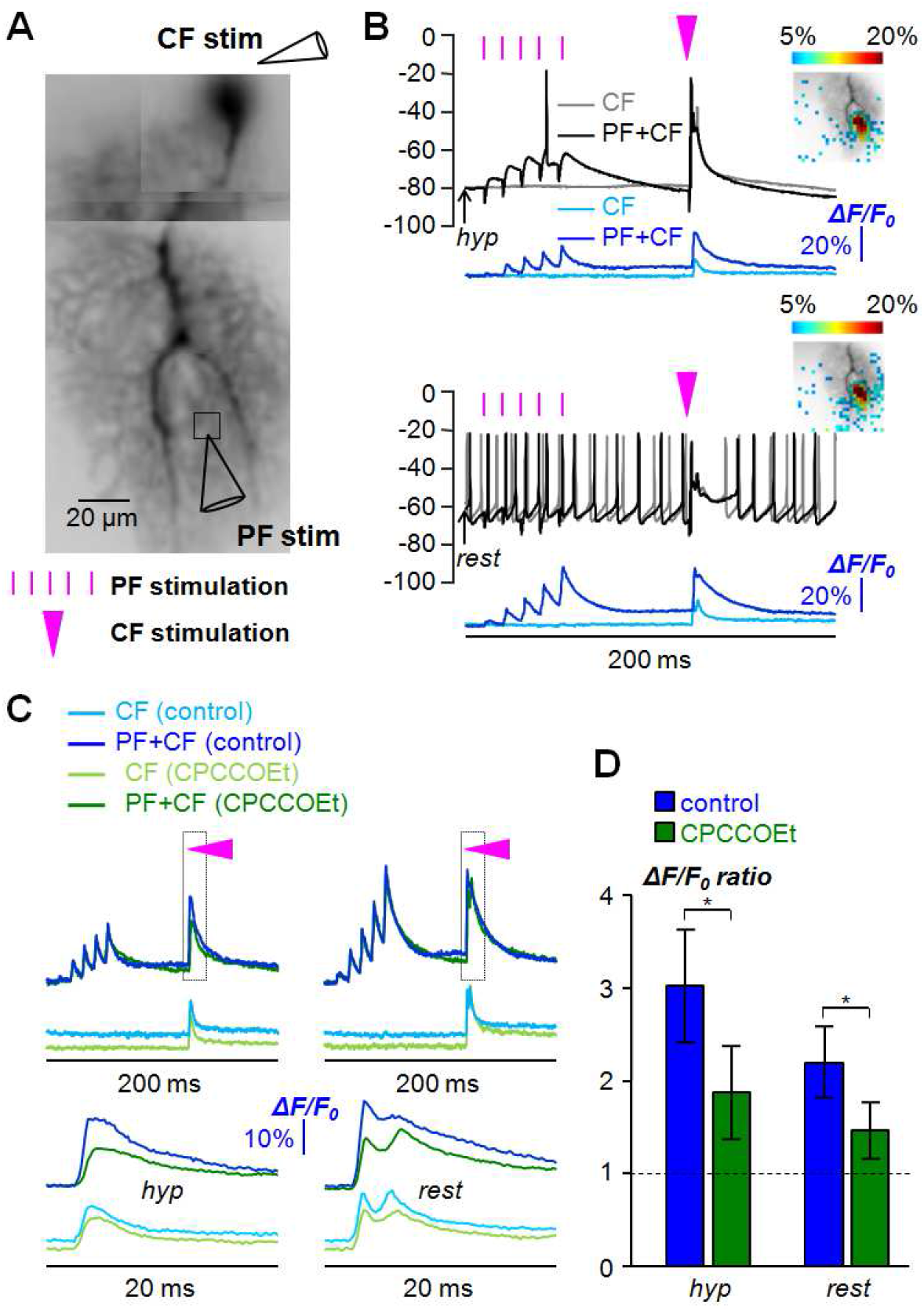
Supralinear Ca^2+^ signals at different initial V_m_. (A) Reconstruction of PN filled with 2 mM OG5N (left) with position of two stimulating electrodes for the CF and PFs and a region of interest adjacent to the PF stimulating electrode indicated. (B) Somatic V_m_ and Ca^2+^ ΔF/F_0_ signals at the region of interest associated with CF stimulation alone (light blue traces) or paired (dark blue traces) with PF stimulation (timing indicated by purple lines) delayed by 110 ms from the first PF pulse (timing indicated by purple triangle) in the case of hyperpolarised (*hyp*, ∼-80 mV) or V_rest_ (*rest*, ∼-60 mV). Spatial distribution of the ΔF/F_0_ signal associated with CF-EPSP paired with PF stimulation depicted using a colour code. (C) Same Ca^2+^ ΔF/F_0_ signals associated with CF stimulation alone or paired with PF stimulation in control solution (blue traces) or after addition of 20 µM of the mGluR1 antagonist CPCCOEt (green traces). The time window outlined is reported below. (D) Mean ± SD (N = 6 cells) of the ΔF/F_0_ ratio between the paired and unpaired signals in control solution or after addition of CPCCOEt. “*” indicates a significant inhibition of supralinear Ca^2+^ signal by CPCCOEt (p< 0.01, paired t-test). The dotted line depicts ΔF/F_0_ ratio = 1.

### Kinetics of dendritic supralinear Ca^2+^ signals

The three forms of supralinear Ca^2+^ signals presented in the previous paragraph are time-correlated with the CF-associated Ca^2+^ transients which are mediated by T-type and P/Q-type VGCCs (Ait Ouares et al., 2019). They can be in principle due to the amplification of these Ca^2+^ sources, to the activation of other Ca^2+^ sources, or to both phenomena. If only the original Ca^2+^ source is amplified, then the kinetics of the rising phase of the Ca^2+^ transient is expected to be the same for the paired and unpaired CF signal. Thus, useful information can be obtained by analysing the kinetics of the rising phase of the Ca^2+^ transients in paired protocols with respect to the Ca^2+^ transients associated with the CF-EPSPs alone, to reveal any possible subsequent component of the Ca^2+^ transients. To this aim, we performed Ca^2+^ recordings associated with the same stimulation protocols described in the previous paragraph, but in this case at 20 kHz to accurately analyse the kinetics of Ca^2+^ transients (Figure 3A). We then analysed the rising phase of the different Ca^2+^ transients exploiting the short relaxation time of OG5N (Jaafari et al., 2015; Jaafari and Canepari, 2016; Ait Ouares et al., 2016). To quantitatively analyse the fast kinetics of Ca^2+^ transients, we averaged 9 recordings for each stimulating protocol (instead of 4), we aligned the Ca^2+^ transients to the beginning of the CF stimulation (Figure 3B), we normalised the signals to the peak and applied a 20-points Savitzky-Golay smoothing filter (Figure 3B), and we finally calculated the time derivative (Jaafari et al., 2014) as shown in Figure 3C. The time to peak of the time derivative, from the CF stimulus, is an indirect kinetics measurement of the Ca^2+^ source and it is typically longer for T-type VGCCs that principally mediate Ca^2+^ influx at *hyp* states, with respect to P/Q-type VGCCs that principally mediate Ca^2+^ influx at *rest* states. Thus, in the example of Figure 3C, the time difference between the two peaks (*Δt_peak_*) at the *hyp* and the *rest* state, in the case of unpaired CF-EPSPs, was 3 samples (150 µs). We then measured the times to peak in the cases of CF-EPSPs in pairing protocol and compared those with the cases of unpaired CF-EPSP. In the example of Figure 3C, the time to peak of the paired CF signal at the *hyp* state was the same of that of unpaired CF signal at the *hyp* state (*Δt_peak_* = 0). In contrast, the times to peak of the paired CF signal at the *rest* state or with the CF stimulation delayed by 60 ms from the first PF stimulus were the same of that of unpaired CF signal at the *rest* state. We performed this analysis in N = 8 cells and calculated both *Δt_peak_* from the CF signal at *hyp* (Figure 3D, top bar diagram) and at *rest* states (Figure 3D, bottom bar diagram). The time to peak of the paired CF signal at *hyp* states was the same of the unpaired CF signal at *hyp* states and the time to peak of the paired CF signal at *rest* states was the same of the unpaired CF signal at *rest* states. In contrast, the time to peak of the CF signals (paired and unpaired) at *hyp* states was significantly (p<0.01 paired t-test) longer (by 2.75 ± 0.71 samples) from the time to peak of the CF signals (paired and unpaired) at *rest* states. Finally, the time to peak of the CF signals at *hyp* states was significantly (p<0.01 paired t-test) longer (by 2.38 ± 0.75 samples) from the time to peak of the paired CF signals with the CF stimulation delayed by 60 ms from the first PF-stimulus. We therefore concluded that the fast kinetics of Ca^2+^ transients associated with the CF-EPSP is identical in paired and unpaired condition, both at *hyp* and *rest* states, when the CF stimulation is delayed by 110 ms from the first PF stimulus. This conclusion suggests that the principal Ca^2+^ source of the supralinear Ca^2+^ signal with the CF stimulation delayed by 110 ms from the first PF stimulus at *hyp* states is the T-type VGCC. In contrast, the principal Ca^2+^ source of the supralinear Ca^2+^ signals with the CF stimulation delayed by 110 ms from the first PF stimulus at *rest* states, or with the CF stimulation delayed by 60 ms from the first PF stimulus, is likely the P/Q-type VGCC.

**Figure 3.**
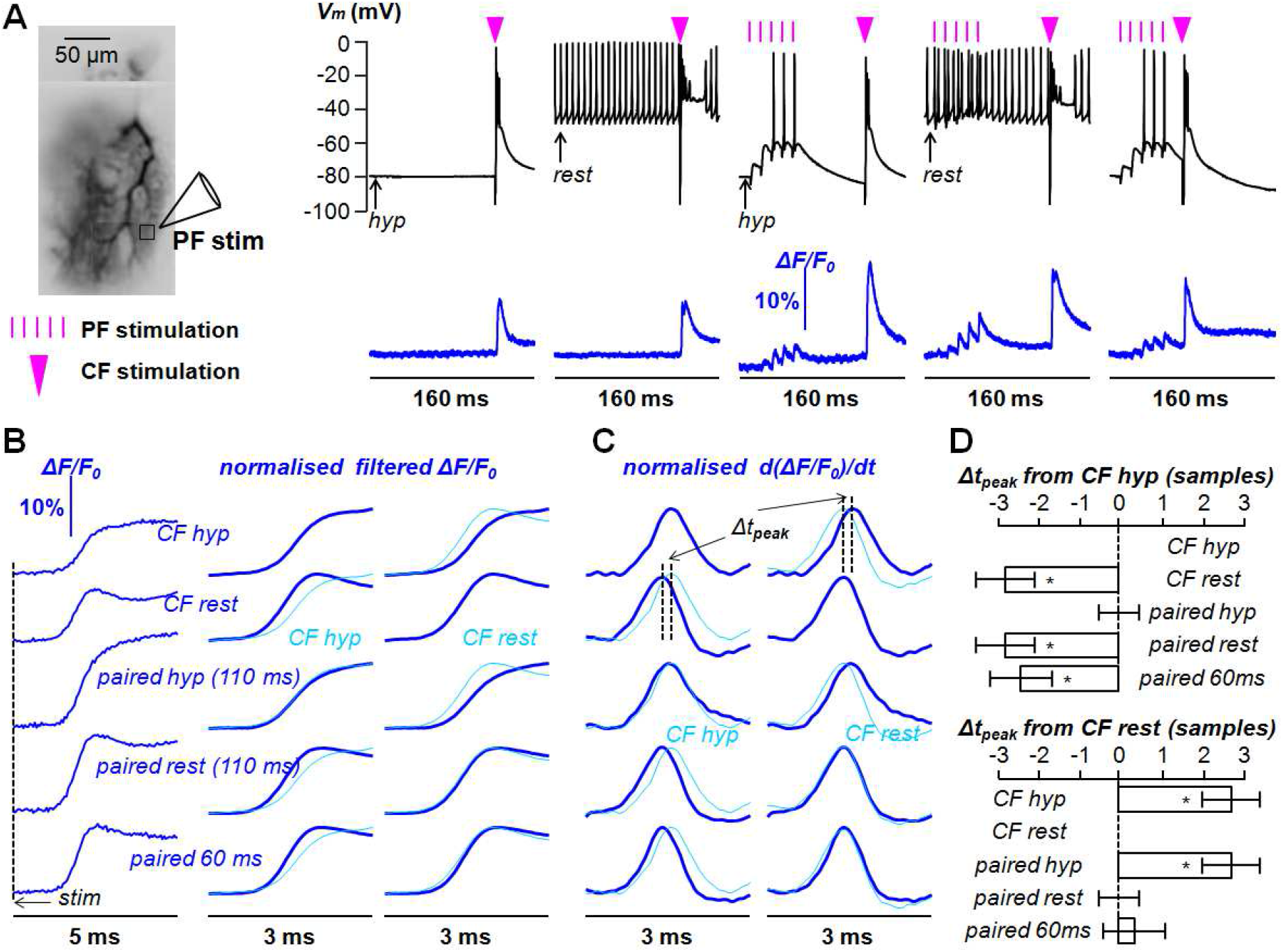
Kinetics of supralinear Ca^2+^ signals in the presence of heparin or ryanodine. (A) Reconstruction of PN filled with 2 mM OG5N; position of PF stimulating electrode and adjacent region of interest indicated. Somatic V_m_ and Ca^2+^ ΔF/F_0_ signals in the region of interest associated with different protocols: from left to right a CF-EPSP alone at *hyp* and *rest* states, paired with a PF-EPSPs train delayed by 110 ms from the first PF stimulus at *hyp* and *rest* states, and paired with a PF-EPSPs train delayed by 60 ms from the first PF stimulus. (B) On the left, Ca^2+^ ΔF/F_0_ signals associated with the CF-EPSP reported in panel A aligned with the CF stimulation (dotted line indicated with “*stim*”) on a time window of 5 ms; on the right, same signals smoothed by a 20-points Savitsky-Golay filter and normalised to the peak on a time window of 3 ms. Traces are reported twice, first superimposed to the signal corresponding to the CF-alone at *hyp* state and second to the signal corresponding to the CF-alone at *rest* state. (C) Normalised time derivative (*d(ΔF/F_0_)/dt*) of the filtered signals in panel B. Traces are reported twice, first superimposed to the time derivative corresponding to the CF-alone at *hyp* state and second to the time derivative corresponding to the CF-alone at *rest* state. The two dotted lines indicate the time of the peak of the *d(ΔF/F_0_)/dt* signals. *Δt_peak_* indicates the time difference between the two peaks for CF-alone at *hyp* state and at *rest* state. (D) Mean ± SD (N = 8 cells) of *Δt_peak_* expressed in samples from the signal associated with the CF-alone at *hyp* state (top) and from the signal associated with the CF-alone at *rest* state (bottom). “*” indicates a significant difference from the signal of reference (p< 0.01, paired t-test).

Yet, several studies have linked PF-triggered activation of mGluR1s to Ca^2+^ release from internal stores, in particular via InsP3 receptors (Finch and Augustine, 2019; Takechi et al., 1998). To directly test whether Ca^2+^ release from internal stores play a role in these supralinear Ca^2+^ signals, we performed experiments with the internal solution containing either 1 mg/mL of heparin (Figure 4A), to inhibit Ca^2+^ store release *via* InsP3 receptors (Kim et al., 2008), or with the internal solution containing 100 µM ryanodine (Figure 4B), to inhibit Ca^2+^ store release *via* ryanodine receptors (Kano et al., 1995). In groups of 6 cells tested with each pharmacological agent, mGluR1-dependent supralinear Ca^2+^ transients were consistent with those measured in control internal solution (Figure 4C), ruling out any significant contribution of Ca^2+^ release from stores either *via* InsP3 or ryanodine receptors.

**Figure 4.**
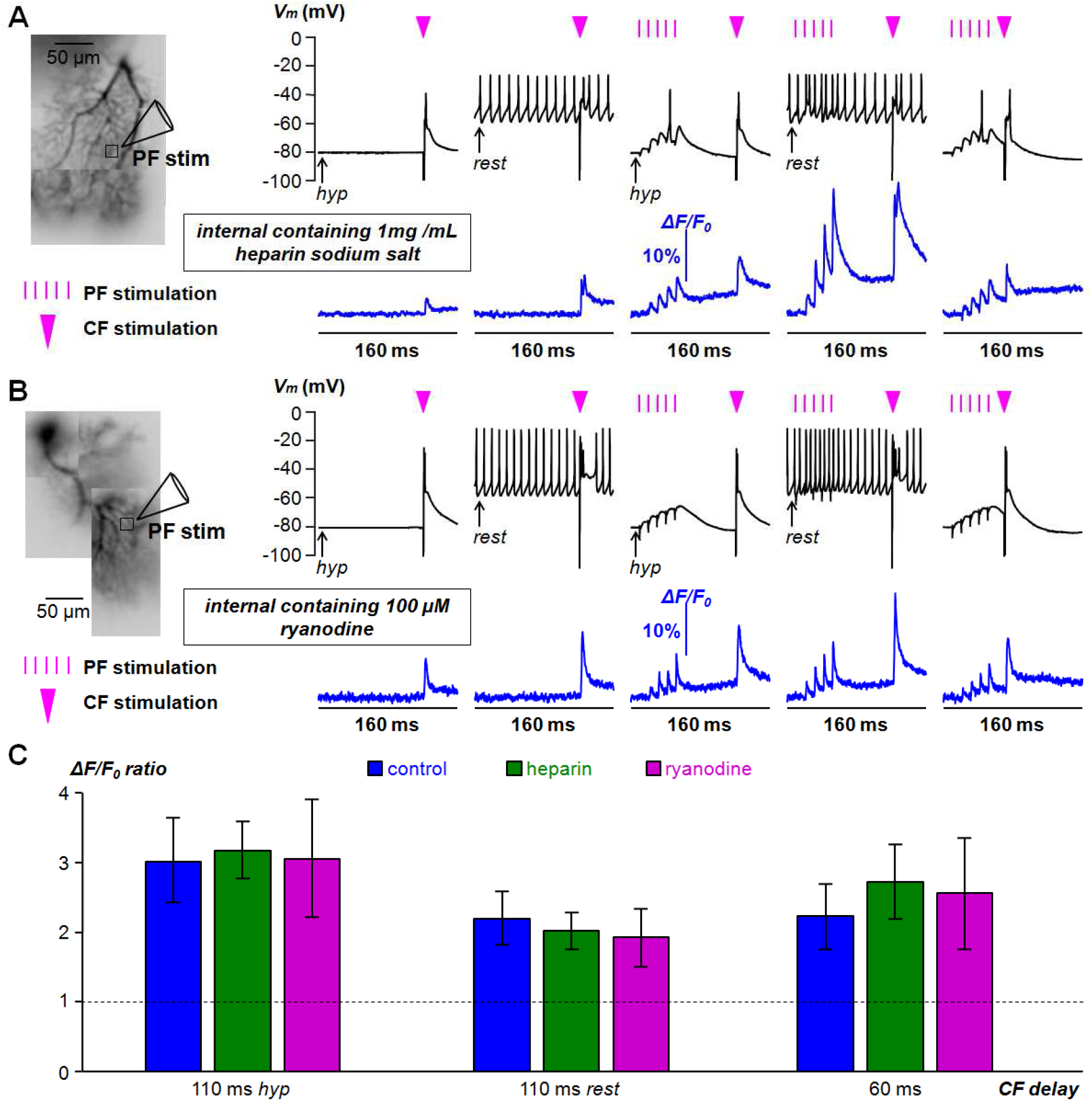
Supralinear Ca^2+^ signals in the presence of heparin or ryanodine. (A) Reconstruction of PN filled with 2 mM OG5N and 1 mg/mL of heparin sodium salt to block InsP3 receptors (left); position of PF stimulating electrode and adjacent region of interest indicated. Somatic V_m_ and Ca^2+^ ΔF/F_0_ signals in the region of interest associated with different protocols: from left to right a CF-EPSP alone at *hyp* and *rest* states, paired with a PF-EPSPs train delayed by 110 ms from the first PF stimulus at *hyp* and *rest* states, and paired with a PF-EPSPs train delayed by 60 ms from the first PF stimulus. (B) Same as A, but with the internal solution containing 100 µM ryanodine to block ryanodine receptors. (C) Mean ± SD (N = 6 cells for each group) of ΔF/F_0_ ratio between the paired and unpaired signals in the presence of heparin (green columns) or ryanodine (purple columns). Blue columns are mean ± SD of ΔF/F_0_ ratios in control conditions already reported in Figure 1 and Figure 2. Values for heparin were: 317 ± 41 for 110 ms CF delay at *hyp* states; 211 ± 48 for 110 ms CF delay at *rest* states; 272 ± 54 for 60 ms CF delay. Values for ryanodine were: 306 ± 85 for 110 ms CF delay at *hyp* states; 193 ± 56 for 110 ms CF delay at *rest* states; 258 ± 80 for 60 ms CF delay. Ryanodine was initially dissolved in DMSO at 25 mM concentration. Experiments were performed 45 minutes after whole cell to obtain full equilibration of the internal solution. Used concentrations were the maximal tolerated by the cells for one-hour recording.

### Dendritic V_m_ associated with mGluR1-dependent supralinear Ca^2+^ signals

Since Ca^2+^ release from internal stores does not contribute to the supralinear Ca^2+^ signals reported above, the origin of these phenomena can be an increase in the Ca^2+^ influx through the plasma membrane during the CF-EPSP, an increase in free Ca^2+^ concentration (and therefore in the Ca^2+^ bound to the OG5N) due to a transient saturation of endogenous Ca^2+^ buffers or a combination of both phenomena. If the supralinear Ca^2+^ signal includes an increase of Ca^2+^ influx through the plasma membrane, the additional charge influx is expected to increase the electrical current and therefore the dendritic V_m_ transient associated with the CF-EPSP. We initially measured dendritic V_m_ associated with a paired CF-EPSP delayed by 60 ms from the first PF stimulus using V_m_ imaging from a region adjacent to the PF stimulation electrode. V_m_ fluorescence transients were calibrated on an absolute scale (in mV) using prolonged hyperpolarising pulses as previously described (Canepari and Vogt, 2008; Ait Ouares et al., 2019), to quantify the depolarisation associated with the CF-EPSP under different conditions. In the experiment reported in Figure 5A, the initial V_m_ before the CF-EPSP shifted from −80 mV to −62 mV when the PF-EPSP train was applied (Figure 5B). Consistently with the fact at that this initial V_m_ A-type K^+^ channels are partially inactivated, and activation of P/Q-type VGCCs is boosted (Ait Ouares et al., 2019), the peak V_m_ shifted from −21 mV to −3 mV. Significant shifts in the initial and peak V_m_ were observed in N = 8 cells in which this experiment was performed (Figure 5C), confirming that the mGluR1-independent supralinear Ca^2+^ signal, occurring when the CF-EPSP is at the end of the PF-EPSPs train, is associated with a shift from T-type VGCCs to P/Q-type VGCCs as suggested by the results reported in Figure 3.

After this first set of V_m_ experiments, we performed combined V_m_ and Ca^2+^ imaging recordings with a similar approach recently utilised to characterise the dendritic V_m_ transient associated with the CF-EPSP alone (Ait Ouares et al., 2019), to measure dendritic V_m_ associated with mGluR1-dependent supralinear Ca^2+^ signals. The cell shown in Figure 6A was filled with the voltage-sentive dye (VSD) JPW1114 and with 1 mM of the low-affinity Ca^2+^ indicator Fura-2FF (K_D_ = 6 µM, Hyrc et al., 2000), used to localise the area of supralinear Ca^2+^ signals both at *hyp* and *rest* initial V_m_ (Figure 6A). Consistently with the observation that PF-ESPSs locally activate VGCCs at *hyp* initial V_m_, the depolarisation associated with the PF train was above −30 mV in the region (*R1*) adjacent to the stimulating electrode, while the depolarisation was comparable with that in the soma in a different region (*R2*), as shown in Figure 6B. To excite VSD fluorescence over 160 ms without causing photodamage, we used only 20% of the laser intensity applied for previous CF-associated V_m_ measurements (Ait Ouares et al., 2019). Using this weaker illumination, the standard deviation of the photon noise was equivalent to ∼2.5 mV after calibration and the peak-to-peak noise was equivalent to ∼10 mV in *R1* (Figure 6B). Therefore, to allow discrimination of V_m_ changes of less than 3 mV, we applied the following filtering procedures (Figure 6C). For the unpaired CF recordings, the initial V_m_ was set to the averaged V_m_ signal before CF stimulation. For the paired CF recordings, the initial V_m_ was accurately estimated by applying a 64-points median filter to the original fluorescence signal after the end of the PF stimulation. Finally, for all recordings, a 4-points median filter was applied to the 20 ms fluorescence signal following CF stimulation. Figure 6D shows a small but detectable additional depolarisation associated with the paired CF-EPSP both at *hyp* and *rest* initial V_m_. In N = 7 cells, in which we successfully performed combined V_m_ and Ca^2+^ with filtered signals above the photon noise to discriminate V_m_ changes < 3 mV, the additional depolarisation, calculated as the difference between the paired and unpaired CF-associated V_m_ transients was always detectable in the case of *hyp* initial V_m_ and corresponded to a significant increase of 5.0 ± 1.6 mV (p< 0.01, paired t-test). In the case of *rest* initial V_m_, a detectable increase (>3 mV) was observed only in 4/7 cells. The mean ± SD of the V_m_ was 2.5 ± 2.5 mV (N = 7 cells). We concluded that small additional V_m_ transients are always associated with mGluR1-dependent supralinear Ca^2+^ signals at *hyp* initial V_m_, a result consistent with the hypothesis that at least part of the supralinear Ca^2+^ signal under this condition is due to Ca^2+^ influx through the plasma membrane. In the case of *rest* initial V_m_, additional V_m_ transients were occasionally observed, suggesting a smaller and possibly highly variable contribution of Ca^2+^ influx through the plasma membrane.

**Figure 5.**
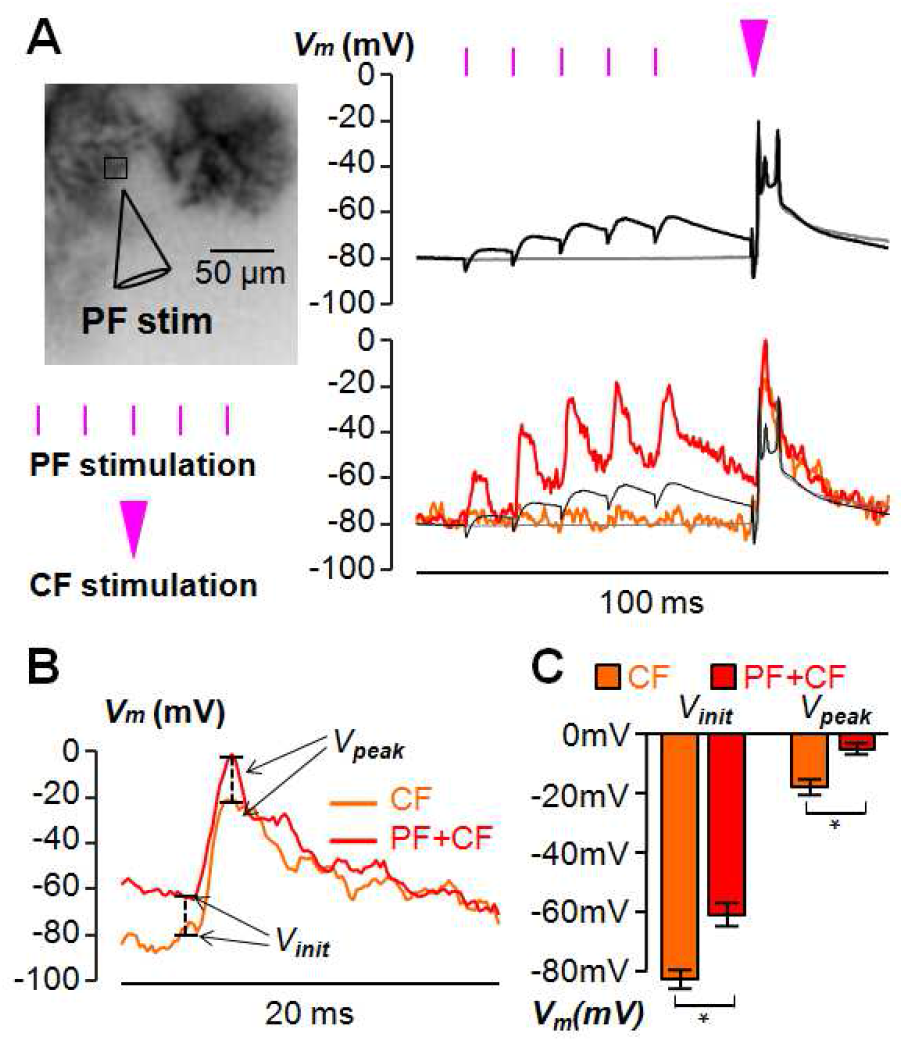
Dendritic V_m_ associated with mGluR1-independent supralinear Ca^2+^ signals. (A) Dendrite of PN filled with 2 mM OG5N (left); position of PF stimulating electrode and adjacent region of interest indicated. Somatic (top) and dendritic (bottom) V_m_ in the region of interest associated with a CF-EPSP alone or paired with a PF-EPSPs train with the CF stimulation delayed by 60 ms from the first PF stimulus. (B) On a time window of 20 ms, dendritic V_m_ signals of panel A. V_init_ and V_peak_ indicate the initial and peak V_m_ calibrated in mV using the procedure previously described in great detail (Ait Ouares et al., 2019). (C) Mean ± SD (N = 8 cells) of V_init_ and V_peak_ for the signals associated with a CF-EPSP alone (−83 ± 3 mV and −18 ± 5 mV respectively) and for the signals associated with a CF-EPSP paired with a PF-EPSPs train with the CF stimulation delayed by 60 ms from the first PF stimulus (−61 ± 4 mV and −5 ± 2 mV respectively). “*” indicates significant differences of V_init_ and V_peak_ (p< 0.01, paired t-test).

**Figure 6.**
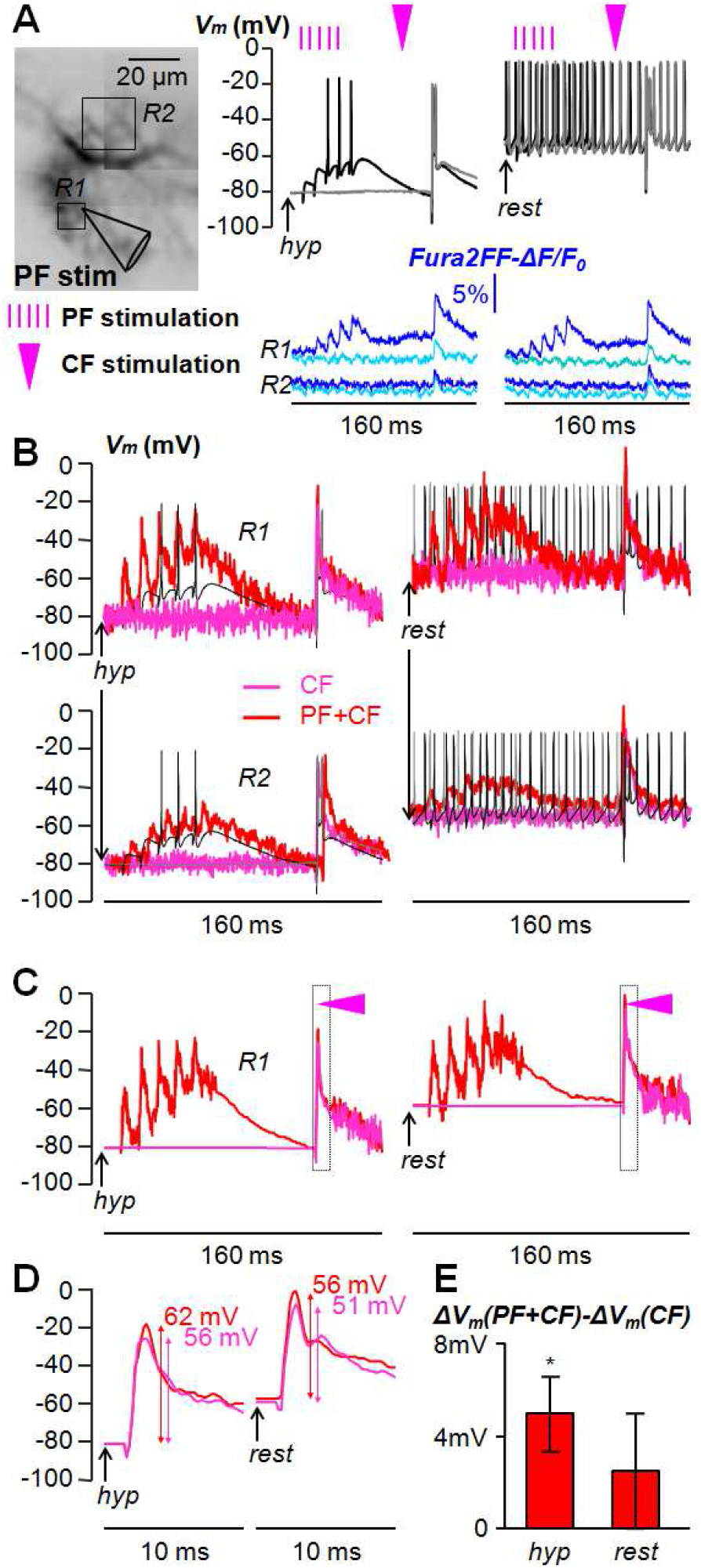
Dendritic V_m_ transients associated with mGluR1-dependent supralinear Ca^2+^ signals. (A) Dendritic PN area filled with the VSD JPW1114 and 1 mM Fura-2FF (left); position of PF stimulating electrode and two regions of interest (*R1* and *R2*) indicated. Somatic V_m_ and Fura-2FF -ΔF/F_0_ signals (right) from single trials at *R1* and *R2* associated with CF stimulation alone (light blue traces) or paired (dark blue traces) with PF stimulation in the case of *hyp* (∼-80 mV) or *rest* (∼-60 mV) initial V_m_, indicating supralinear Ca^2+^ signals at *R1*. (B) V_m_ calibrated recordings associated with CF stimulation alone (red traces) or paired (pink traces) with PF stimulation at *R1* and *R2* superimposed to somatic V_m,_ in the case of *hyp* or *rest* initial V_m_. (C) V_m_ recordings at *R1* shown in panel B filtered as following: for the CF alone, average of the signals before the CF stimulation and in the next 20 ms a 4-points median filter is applied; for the paired CF, a 64-points median filter is applied after the PF train and before the CF stimulation and in the next 20 ms a 4-points median filter is applied. (D) Same signals of panel C in the outlined time window; the values of the V_m_ transients are indicated. (E) Mean ± SD (N = 7 cells) of the V_m_ transient difference between the paired and unpaired CF in the case of *hyp* or *rest* initial V_m_. “*” indicates a significant difference (p< 0.01, paired t-test).

### Contribution of the saturation of the endogenous buffer to the supralinear Ca^2+^ signals

In a previous study we have shown that trains of PF stimuli at 100 Hz produce a local progressive saturation of endogenous Ca^2+^ buffers that boosts a Ca^2+^ fluorescent transient associated with a CF-EPSP occurring at the end of the PF train (Canepari and Vogt, 2008). Since mGluR1 activation induces a slow Ca^2+^ influx, saturation of the endogenous Ca^2+^ buffer may occur on a longer time scale and contribute to the mGluR1-dependent supralinear Ca^2+^ signals. To test whether this phenomenon occurs, we performed experiments including the high-affinity UV-excitable indicator Fura-2 (Fura2, K_D_ ∼200 nM), at the concentration of 400 µM. The fast mobile endogenous Ca^2+^ buffer that dominates Ca^2+^ binding in the millisecond time scale is Calbindin D-28k (Schmidt and Eilers, 2009), a protein with four binding sites and an estimated concentration of 120 µM in PN dendrites (Kosaka et al., 1993). The two pairs of binding sites have estimated k_ON_ of 43.5 µM^−1^ s^−1^ and 5.5 µM^−1^ s^−1^ respectively, and estimated k_D_ of 822 nM and 474 nM respectively (Nägerl et al., 2000). Since the association constant of Fura2 is faster than that of proteins forming the endogenous buffer (k_ON_^Fura2^ ∼ 500 µM^−1^ s^−1^, see Canepari and Mammano, 1999; Ait Ouares et al., 2019), Ca^2+^ binding to Fura2 will anticipate Ca^2+^ binding to the endogenous buffer at very short time scale. In addition, since the total concentration of Fura2 (400 µM) is close to the expected binding sites concentration of Calbindin D-28k (480 µM), the fraction of Fura2 bound to Ca^2+^ will be comparable to that binding to Calbindin D-28k in the absence of Fura2. Thus, the analysis of the saturation of Fura2 will provide an estimate for the saturation of the endogenous buffer occurring physiologically. In the cell of Figure 7A, we applied the same stimulation protocols of Figure 3 and measured separately the absolute fluorescent transients from the two indicators (OG5N ΔF/F_0_ and Fura2 -ΔF/F_0_). Under this condition, regular supralinear OG5N Ca^2+^ signals were observed (Figure 7B), indicating that these signals are not prevented by the presence of Fura2 at this moderate concentration. In contrast, whereas a supralinear Fura2 Ca^2+^ signal associated with the paired CF-EPSP delayed by 110 ms at the *hyp* state was observed, the Fura2 Ca^2+^ signal associated with the paired CF-EPSP delayed by 60 ms was smaller than the Ca^2+^ signal associated with the unpaired CF-EPSP (Figure 7B). To obtain a quantitative interpretation of this qualitative observation, we calculated the variable “*S*” defined as the ratio of the ΔF/F_0_ peaks for each protocol against the ΔF/F_0_ peak associated with the unpaired CF stimulation (Figure 7C). The conclusive information that can be obtained, by the analysis of the variable *S*, is explained in detail in the *Experimental design and statistical analysis* section of the Materials and Methods. Briefly, if a supralinear Ca^2+^ signal is exclusively due to buffer saturation, then the value of *S* for Fura2 (*S_Fura2_*) must be less than one. Thus, a value of *S_Fura2_* larger than 1 indicates that larger Ca^2+^ influx contributes to the supralinear Ca^2+^ signal. If, in contrast, a supralinear Ca^2+^ signal is exclusively due to larger Ca^2+^ influx, (i.e. no buffer saturation), then the increase in the absolute ΔF/F_0_ signal will be the same for the two indicators. Thus, in this case, the ratio of *S* for the two indicators (*S_Fura2_*/*S_OG5N_*) must be 1 while a significantly lower value of this ratio indicates that buffer saturation contributes to the supralinear Ca^2+^ signal. Finally, if two supralinear Ca^2+^ signals are calculated from the same unpaired CF signal, i.e. from the same denominator at *hyp* states, then *S_Fura2_*/*S_OG5N_* represents a quantitative estimate of the extent of buffer saturation in the two cases. In N = 8 cells tested, *S_Fura2_* was significantly positive for the mGluR1-dependent supralinear Ca^2+^ signal at *hyp* states (p<0.01 paired t-test, *S_Fura2_* = 2.38 ± 0.05), confirming that larger Ca^2+^ influx through the plasma membrane contributes in this case. However, *S_Fura2_* was significantly smaller than *S_OG5N_* (p<0.01, paired t-test, *S_Fura2_*/*S_OG5N_* = 0.58 ± 0.09) indicating that also buffer saturation contributes to this supralinear Ca^2+^ signal. Finally, *S_Fura2_*/*S_OG5N_* for the mGluR1 independent supralinear Ca^2+^ signal (0.31 ± 0.13) was significantly smaller than *S_Fura2_*/*S_OG5N_* for the mGluR1-dependent supralinear Ca^2+^ signal (p<0.01, paired t-test), indicating that in this case buffer saturation provides a larger contribution. The Ca^2+^ increase through P/Q-type VGCCs produced by switching from *hyp* states to *rest* states is essentially due to larger Ca^2+^ influx (*S_Fura2_*/*S_OG5N_* = 0.80 ± 0.23). Thus, as expected, the further mGluR1-dependent Ca^2+^ increase includes a significant contribution of buffer saturation (*S_Fura2_*/*S_OG5N_* = 0.54 ± 0.19, calculated using the unpaired CF signal at *rest* states). This analysis, however, does not allow estimating a possible contribution of larger Ca^2+^ influx in the case of mGluR1-dependent supralinear Ca^2+^ signal at *rest* states.

**Figure 7.**
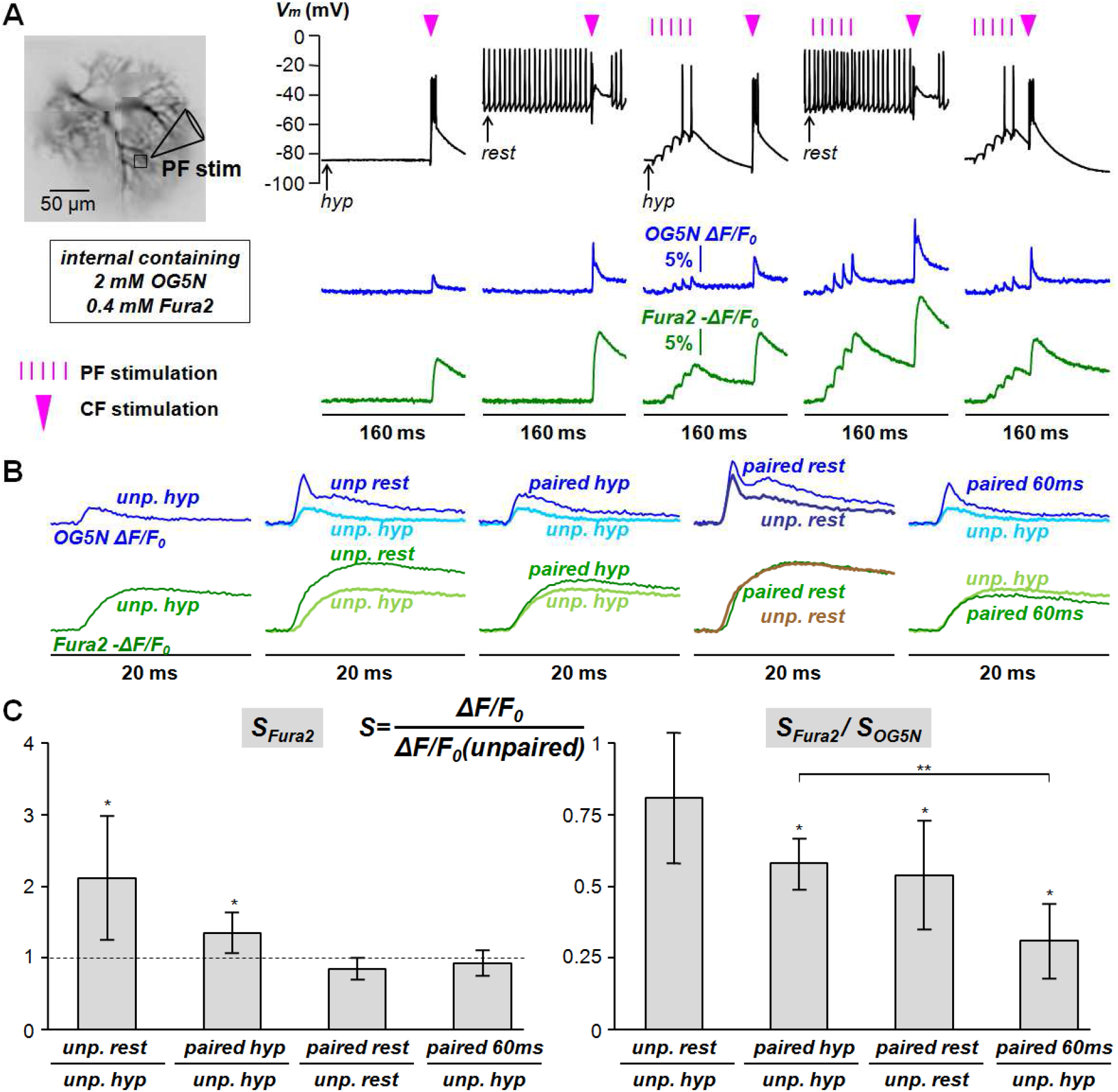
Contribution of saturation of endogenous Ca^2+^ buffers to supralinear Ca^2+^ signals. (A) Dendritic PN area filled with two Ca^2+^ indicators (left): OG5N (2 mM, K_D_ = 35 µM) and Fura2 (0.4 mM, K_D_ = 0.2 µM); position of PF stimulating electrode and adjacent region of interest indicated. Somatic V_m_ and Ca^2+^ signals in the region of interest associated with different protocols: from left to right a CF-EPSP alone at *hyp* and *rest* states, paired with a PF-EPSPs train delayed by 110 ms from the first PF stimulus at *hyp* and *rest* states, and paired with a PF-EPSPs train delayed by 60 ms from the first PF stimulus; signals from the two indicators (OG5N ΔF/F_0_ and Fura -ΔF/F_0_) were obtained by alternating the excitation wavelength. (B) Same Ca^2+^ transients associated with the CF-EPSP in panel A, for the two indicators, superimposed either to the unpaired CF transient at *hyp* state or to the unpaired CF transient at *rest* state. (C) *S* is defined as the ratio between the ΔF/F_0_ peaks for each protocol and the ΔF/F_0_ peak associated with the unpaired CF stimulation. On the left, mean ± SD (N = 8 cells) of *S* corresponding to Fura2 signals (*S_Fura2_*) in the different cases as reported in the legend; “*” indicates a significant difference (p< 0.01, paired t-test) between the two ΔF/F_0_ peaks used to calculate *S_Fura2_*. On the right, mean ± SD (N = 8 cells) of *S_Fura2_*/*S_OG5N_* in the different cases as reported in the legend; “*” indicates a significant difference (p< 0.01, paired t-test) between *S_OG5N_* and *S_Fura2_*. “**” indicates a significant difference (p< 0.01, paired t-test) between two *S_OG5N_*/*S_Fura2_* ratios.

### Origin of the mGluR1-dependent supralinear Ca^2+^ signals at initial hyperpolarised V_m_

The results presented above demonstrate that mGluR-dependent supralinear Ca^2+^ signals at the *hyp* state are due to a combination of Ca^2+^ influx increase through the plasma membrane and to a transient saturation of the endogenous Ca^2+^ buffer. The rising phase of the paired CF Ca^2+^ signal has the same kinetics of the Ca^2+^ signal associated with the CF alone (Figure 3) which is principally mediated by T-type VGCCs (Ait Ouares et al., 2019). It has been reported that mGluR1 activation potentiate Cav3.1 T-type VGCCs in PNs (Hildebrand et al., 2009). Hence, we finally assessed directly whether the Ca^2+^ influx component of the mGluR1-dependent supralinear Ca^2+^ signal at *hyp* states is through these channels. In the cell reported in Figure 8A, we recorded the Ca^2+^ transient associated with a CF-EPSP alone or paired with PF stimulation, at *hyp* or *rest* initial V_m_ in control conditions with a delay of 110 ms from the first PF stimulus and the CF stimulation. Addition of 30 µM of the Cav3.1 blocker NNC550396 (NNC) inhibited spontaneous firing at *rest* initial V_m_ and strongly reduced Ca^2+^ transients associated both with PF-EPSPs and the CF-EPSP at *hyp* initial V_m_ (Figure 8A). In contrast, Ca^2+^ transients associated both with PF-EPSPs and the CF-EPSP at *rest* initial V_m_ were only slightly affected by NNC addition, confirming that dendritic T-type VGCCs are mostly inactivated at this state (Figure 8A). As shown in Figure 8B, the supralinear Ca^2+^ transient at *hyp* initial V_m_ was also strongly inhibited by addition of NNC. In N = 6 cells tested with this experimental protocol, the ratios between the CF Ca^2+^ signals in the presence of NNC and in control condition at initial hyperpolarised V_m_ were 0.44 ± 0.18 and 0.32 ± 0.13 for the CF alone or paired with PF stimulation respectively (Figure 8C). In both cases, the Ca^2+^ signal associated with the paired CF-EPSP was significantly smaller (p< 0.01, paired t-test). In contrast, the ratios between the CF Ca^2+^ signals in the presence of NNC and in control condition at *rest* initial V_m_ were 0.93 ± 0.13 and 0.91 ± 0.10 for the CF alone or paired with PF stimulation respectively, indicating that in this state T-type VGCCs contribute marginally to Ca^2+^ signals (Figure 8C). The results reported in Figure 8 unambiguously confirm that the Ca^2+^ influx component responsible for mGluR1-dependent supralinear Ca^2+^ signals at *hyp* state is mediated by T-type Ca^2+^ channels.

**Figure 8.**
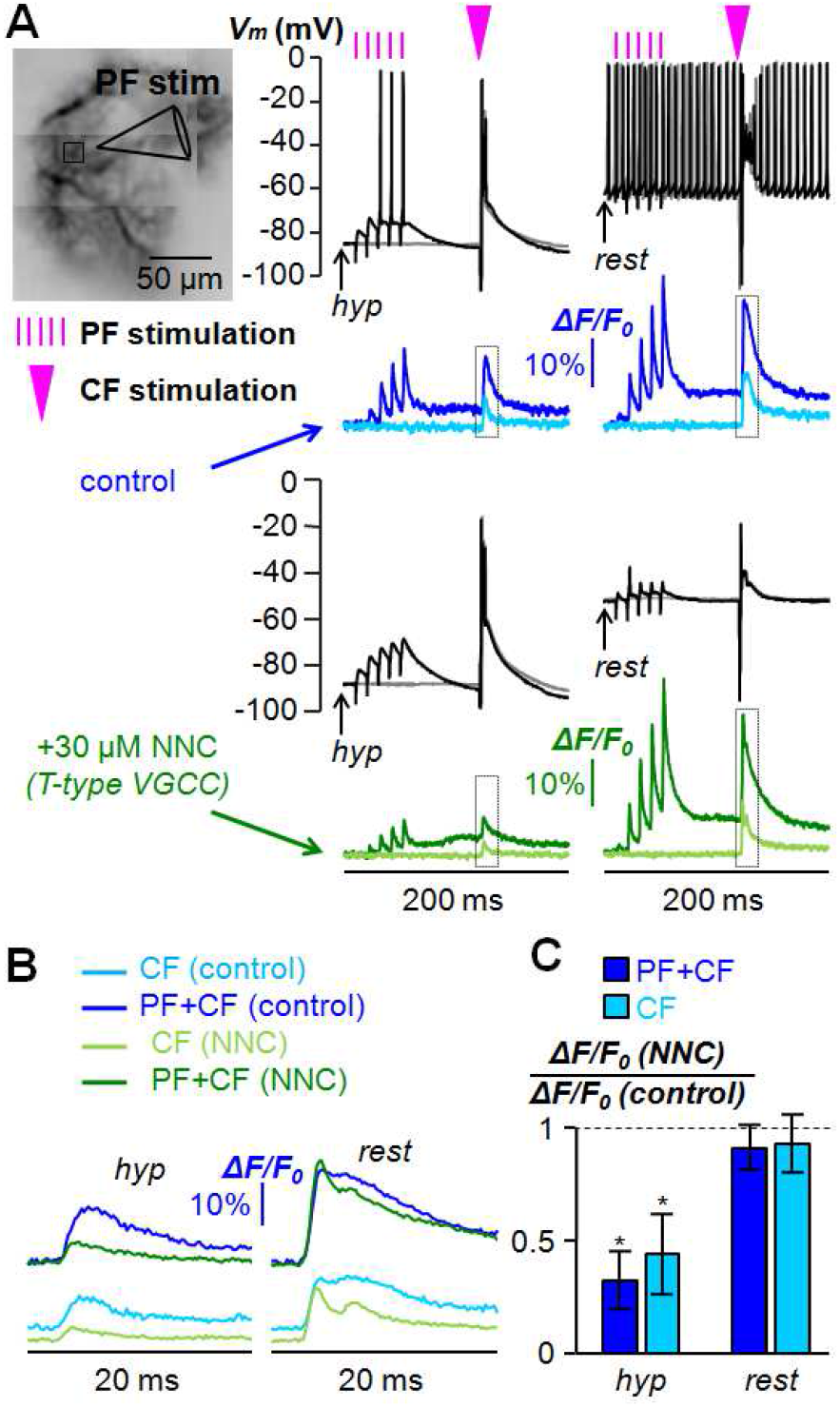
Supralinear Ca^2+^ signals after blocking T-type VGCCs. (A) Dendritic area of a PN filled with 2 mM OG5N (left) with position of PF stimulating electrode and an adjacent region of interest indicated. On the right, somatic V_m_ (black traces) and Ca^2+^ ΔF/F_0_ (blue or green traces) signals at the region of interest associated with CF stimulation alone (light traces) or paired (dark traces) with PF stimulation (timing indicated by purple lines) delayed by 110 ms from the first PF pulse (timing indicated by purple triangle) in the case of *hyp* or *rest* initial V_m_. Signals were acquired in control conditions (blue traces) or after addition of 30 µM of the T-type VGCC blocker NNC (green traces). (B) Same Ca^2+^ signals in panel A over a 20 ms time window outlined. (C) Mean ± SD (N = 6 cells) of the ratios between the ΔF/F_0_ peak after addition of NNC and the peak under control condition; dark blue columns are for paired signals; light blue columns are for CF alone signals. Ratios are calculated both for unpaired and unpaired CF signals. “*” indicates a significant reduction of paired Ca^2+^ signal after addition of NNC (p< 0.01, paired t-test). The dotted line depicts the ratio = 1.

As shown in the example of Figure 8A, addition of NNC reduced the fast Ca^2+^ transient associated with the PF-EPSP train while it did not affect the slow fast Ca^2+^ transient mediated by mGluR1s. Furthermore, although both unpaired and paired Ca^2+^ transients were inhibited by NNC, the paired Ca^2+^ transient was still larger (Figure 8A), suggesting that local amplification of the residual component by buffer saturation is still present when T-type VGCCs are blocked. Thus, endogenous buffer is likely saturated by the slow mGluR1-mediated Ca^2+^ influx. To test this hypothesis, we measured mGluR1-dependent supralinear Ca^2+^ signals at *hyp* states while blocking the slow mGluR1-mediated Ca^2+^ transient with the channel blocker IEM1460 that does not affect the mGluR1-dependent boosting of T-type VGCCs (Hildebrand et al., 2009). In the example of Figure 9A, addition of 250 µM of IEM1460 produced a use dependent block of the slow mGluR1-dependent Ca^2+^ transient over a period of 20 minutes in which stimulation protocols were repeated every 5 minutes. While a change in the somatic V_m_ recordings was observed after IEM1460 application, the fast PF-mediated Ca^2+^ transients did not change indicating that VGCCs were not directly affected by the blocker. Whereas the Ca^2+^ transient associated with the unpaired CF-EPSP was also not affected, the Ca^2+^ transient associated with the paired CF-EPSP was progressively reduced consistently with the slow mGluR1-mediated Ca^2+^ transient that was extrapolated using a simple fitting of the decay phase of the fast PF-mediated Ca^2+^ transient (Figure 9B). We performed this experiment in N = 6 cells, obtaining a significant reduction of the slow mGluR1-mediated Ca^2+^ transient (p<0.01, paired t-test), 20 minutes after IEM1460 application, corresponding to a ΔF/F_0_ ratio of 0.49 ± 0.16 (Figure 9C). Consistently with this result, the supralinear Ca^2+^ signal, quantified again as the ratio between the unpaired and the paired ΔF/F_0_ peaks, significantly (p < 0.01, paired t-test) decreased from 3.37 ± 0.83 to 1.86 ± 0.56 (Figure 9D). We concluded that the mGluR1-dependent supralinear Ca^2+^ signal at *hyp* states are caused by boosting of T-type VGCCs amplified by endogenous buffer saturation produced by the slow mGluR1-dependent PF-mediated Ca^2+^ influx.

**Figure 9.**
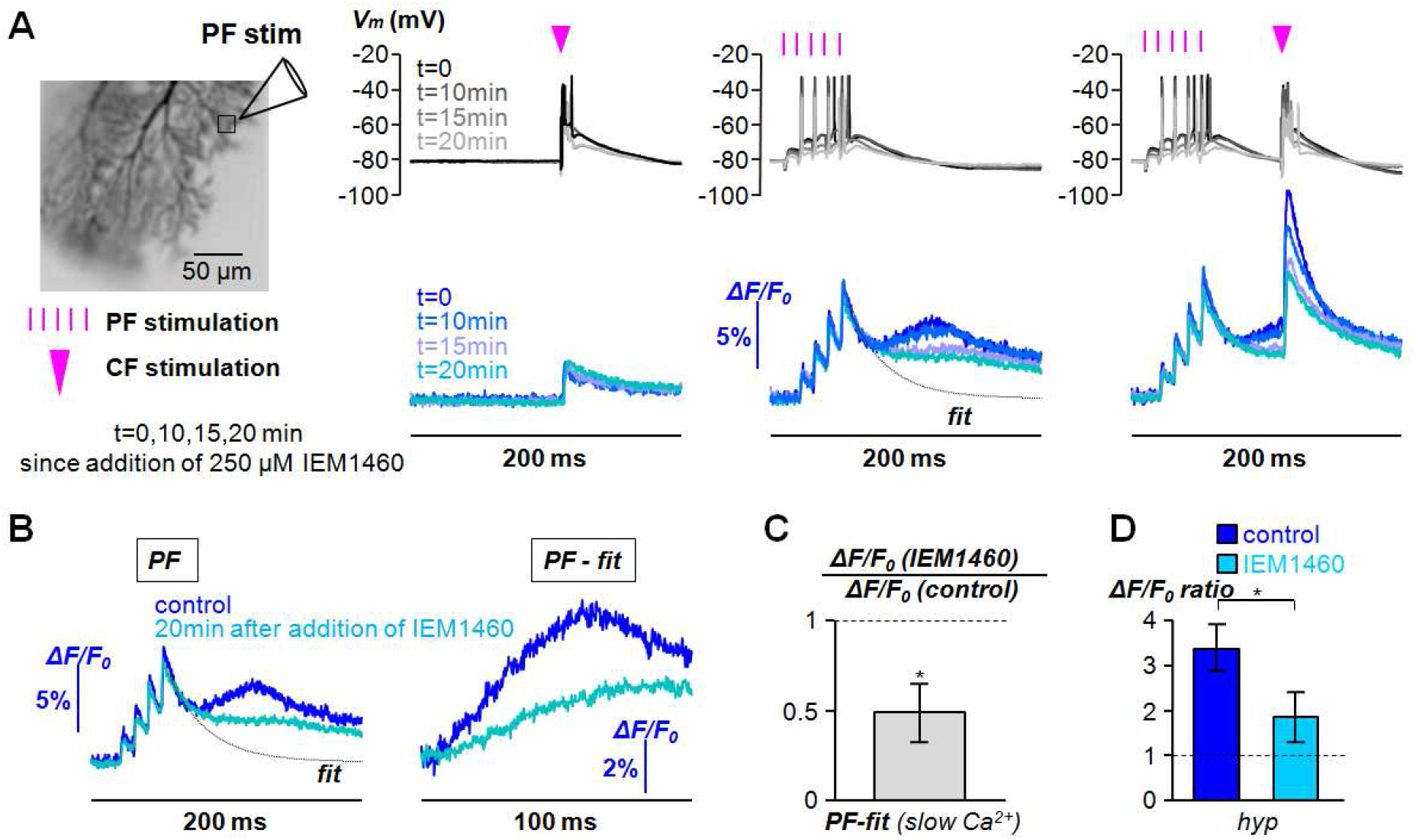
Use dependent block of mGluR1 slow Ca^2+^ transients and of supralinear Ca^2+^ signals. (A) Dendritic area of a PN filled with 2 mM OG5N (left) with position of PF stimulating electrode and an adjacent region of interest indicated. Somatic V_m_ and Ca^2+^ ΔF/F_0_ signals at the region of interest associated with a CF-EPSP alone, with PFs stimulated with 5 pulses at 100 Hz or with a CF-EPSP paired with PF stimulation at *hyp* state. Black and dark blue traces were from recordings in control solution. Traces at different gray or blue tones were from recordings performed a time (t) after addition of 250 µM of the channel blocker IEM1460. (B) On the left, same recordings in A for PF stimulation only (*PF*) in control condition and 20 minutes after addition of IEM1460; the exponential “*fit*“ of the Ca^2+^ ΔF/F_0_ decay after the 5^th^ PF-EPSP is also reported with a dotted line. On the right, Ca^2+^ transients associated with the slow mGluR1-mediated component calculated as the difference between the Ca^2+^ ΔF/F_0_ signals associated with PF stimulation and the *fit* trace on the left. (C) Mean ± SD (N = 6 cells) of the ratio between the slow mGluR1-dependent PF-mediated Ca^2+^ transients (*PF -fit*) 20 minutes after addition of IEM1460 and in control condition. “*” indicates a significant reduction of the signal (p< 0.01, paired t-test). Dotted line depicts the ratio = 1. (D) Mean ± SD of the ΔF/F_0_ ratio between the paired and unpaired signals in control solution or after addition of IEM1460. “*” indicates a significant inhibition of supralinear Ca^2+^ signal by CPCCOEt (p< 0.01, paired t-test). The dotted line depicts ΔF/F_0_ ratio = 1.

### Origin of the mGluR1-dependent supralinear Ca^2+^ signals at V_rest_

The experiments reported in Figure 9 show that the mGluR1-dependent supralinear Ca^2+^ signal at *rest* state is largely due to saturation of the endogenous buffer, but the results reported in Figure 8 also suggest that an increase in Ca^2+^ influx may contribute in a variable manner. It has been reported that mGluR1 activation shifts the inactivation curve of A-type K^+^ channels boosting the opening of P/Q VGCCs at initial V_m_ > −65 mV in PNs (Otsu et al., 2014). Thus, consistently with the results of the kinetics analysis reported in Figure 3, an increase in Ca^2+^ influx *via* P/Q-type VGCCs may contribute to this supralinear Ca^2+^ signal. In contrast to the direct local regulation of T-type VGCCs by mGluR1s (Hildebrand et al., 2009), occurring at *hyp* states, the indirect regulation of P/Q-type VGCCs is produced by a V_m_ modulation and therefore it is not co-localised with mGluR1s. By analysing cells in which V_rest_ was <-60 mV, i.e. where A-type K^+^ channels are not yet fully inactivated, we observed some occasional small supralinear Ca^2+^ signals at far-away sites from the PF-activated region at *rest* states. An example of this wide supralinear Ca^2+^ signal is reported in Figure 10A-B. While this far supralinear Ca^2+^ signal was smaller than the supralinear Ca^2+^ signal co-localised with the PF-mediated signal (Figure 10A), the kinetics was qualitatively similar to that of a “supralinear” Ca^2+^ signal obtained by blocking A-type K^+^ channels with the toxin AmmTx3 (Zoukimian et al., 2019; Ait Ouares et al., 2019), as shown in the example of Figure 10C.

**Figure 10.**
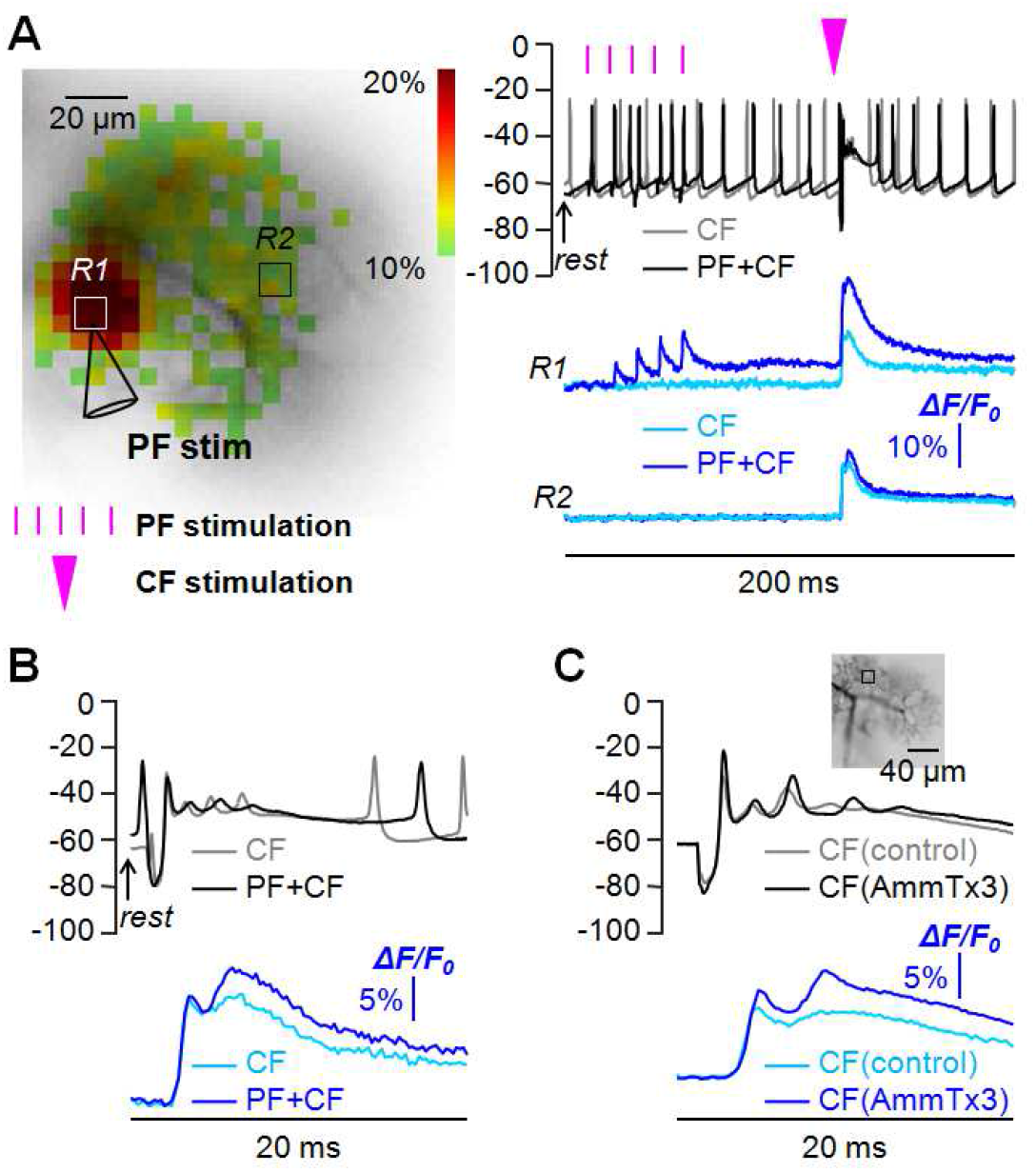
Wide mGluR1-dependent supralinear Ca^2+^ signals at *rest* states. (A) On the left, dendrite of PN filled with 2 mM OG5N; position of PF stimulating electrode and of two regions of interest indicated (*R1* and *R2*); *R1* is adjacent to the PF stimulating electrode; R2 is ∼60 µm away from the PF stimulating electrode. On the right, somatic V_m_ (top) and dendritic Ca^2+^ signals (bottom) in *R1* and *R2* associated with a CF-EPSP alone or paired with a PF-EPSPs train with the CF stimulation delayed by 110 ms from the first PF stimulus at *rest* state. (B) On a time window of 20 ms, somatic V_m_ and dendritic Ca^2+^ signals in *R2* reported in panel A, exhibiting a supralinear Ca^2+^ transient smaller than that in *R1*. (C) Somatic V_m_ (top) and dendritic Ca^2+^ signal associated with a CF-EPSP alone in control condition or after addition of the A-type K^+^ channel inhibitor AmmTx3. The two supralinear Ca^2+^ signals reported in panels A and B are qualitatively very similar.

Finally, in many studies, physiological mGluR1 activation has been mimicked by bath application of the agonist (S)-3,5-dihydroxyphenylglycine (DHPG, see for example Canepari et al., 2001; Tempia et al., 2001; Maejima et al., 2005; Otsu et al., 2014). We examined this artificial stimulation protocol by applying 100 µM DHPG, using a pipette positioned near the dendrites, a short (20 ms) pressure ejection as shown in the example of Figure 12A. At *hyp* state, DHPG application triggered a slow Ca^2+^ signal, but in contrast to the physiological PF stimulation a corresponding slow V_m_ depolarisation was observed in the soma (Figure 12A). Thus, while a substantial supralinear Ca^2+^ signal associated with DHPG pairing was observed, the kinetics of the Ca^2+^ transient indicates a dominance of Ca^2+^ influx via P/Q-type VGCCs (Figure 12B), consistently with the fact that the initial V_m_ for the CF-EPSP is ∼-60 mV. The size of the supralinear Ca^2+^ signal associated with DHPG application is comparable to those associated with PF stimulation (Figure 12C). Nevertheless, since all these phenomena are largely due to transient saturation of the endogenous buffer and the spatial and temporal dynamics of Ca^2+^ influx causing this saturation is different in the different protocols, we conclude that experiments using DHPG application cannot realistically mimic the physiological scenarios associated with mGluR1 activation.

**Figure 11.**
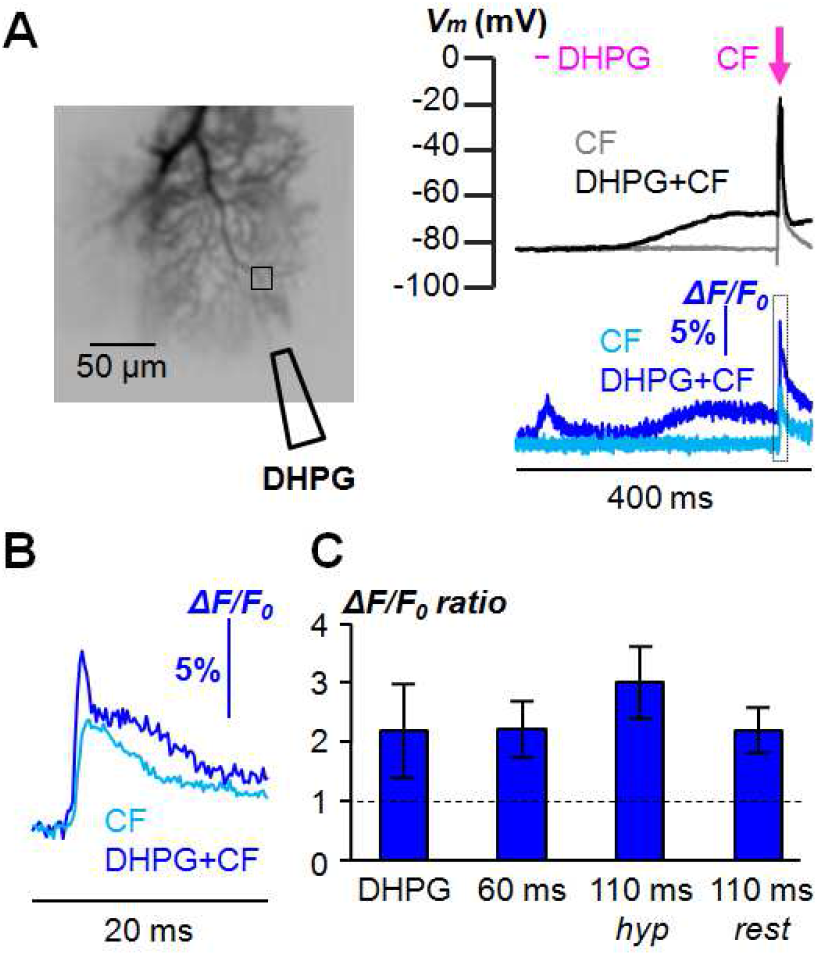
Dendritic V_m_ associated with mGluR1-independent supralinear Ca2+ signals. (A) On the left, dendrite of PN filled with 2 mM OG5N (left); a region of interest and the position of a pipette delivering 100 µM DHPG dissolved in external solution are indicated; DHPG application was achieved by applying a 20 ms pulse of pressure using a pressure ejector PDES-2DX (npi, Tamm, Germany). On the right, somatic V_m_ (top) and dendritic Ca^2+^ signals (bottom) in the region of interest associated with a CF-EPSP alone (light blue trace) or paired with a DHPG application (dark blue trace); the timing of DHPG application and of CF stimulation are indicated with a purple line and a purple arrow respectively; a ΔF/F_0_ artefact is observable during the short DHPG application. (B) On a time window of 20 ms, dendritic Ca^2+^ signals reported in panel A, exhibiting a supralinear Ca^2+^ transient. (C) First on the left, mean ± SD (2.20 ± 0.80, N = 7 cells) of ΔF/F_0_ ratio between the CF-associated signal paired with DHPG application and alone. Following on the right, mean ± SD of ΔF/F_0_ ratio between CF-associated signal paired with PF stimulation and alone; the statistics relative to 60 ms delay between the first PF stimulus and the CF stimulation is the same reported in Figure 1D; the two statistics relative to 110 ms delay between the first PF stimulus and the CF stimulation at *hyp* and *rest* states are the same reported in Figure 2D. The dotted line depicts ΔF/F_0_ ratio = 1.

**Figure 12.**
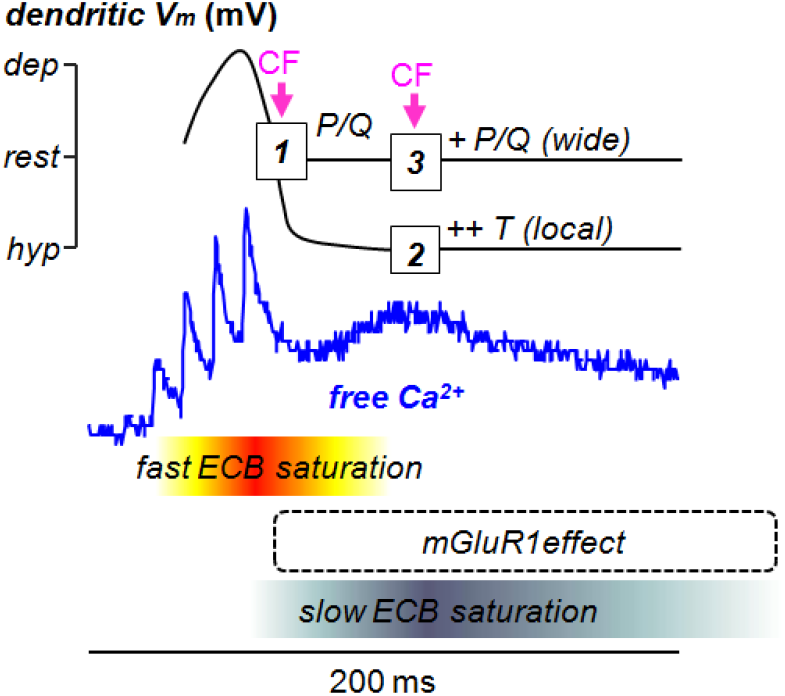
Determinants of different dendritic supralinear Ca^2+^ signals in PNs. A representative transient of free Ca^2+^ concentration associated with of 5 PF-EPSPs at 100 Hz (derived from a cell analysed in this study) is depicted (blue trace). The dendritic V_m_ at the PF site is transiently depolarised during the fast PF-EPSPs. Thus, when a CF-EPSP occurs during this first time window (box *1*), it activates P/Q-type VGCCs. The dendritic V_m_ after the fast PF-EPSPs is repolarised while the effects of mGluR1s become active. In this second time window, if the dendritic V_m_ is at the *hyp* state, a CF-EPSP activates T-type VGCCs which are strongly (++) and locally potentiated by an mGluR1 action (box *2*). In contrast, if the dendritic V_m_ is at the *rest* state, a CF-EPSP activates P/Q-type VGCCs which are indirectly, variably (+) and widely potentiated by an mGluR1 action to A-type K^+^ channels. All free Ca^2+^ concentration transients associated with a CF-EPSP paired with PF-EPSPs are locally larger than those associated with a CF-EPSP alone. This supralinear Ca^2+^ signal component is due to the transient saturation of endogenous Ca^2+^ buffer (ECB) initially by Ca^2+^ influx via VGCCs (fast ECB saturation) and later by the slow mGluR1-dependent Ca^2+^ influx (slow ECB saturation).

In summary, at the *rest* state, regulation of P/Q-type VGCCs by mGluR1 activation may contribute to the supralinear Ca^2+^ signal, but the localisation of this signal, produced by Ca^2+^ influx *via* these channels, is entirely due to the previous Ca^2+^ entry that saturates the endogenous buffer.

## DISCUSSION

### Determinants of the different dendritic supralinear Ca^2+^ signals in PNs

In this study we identified the biophysical determinants of the dendritic supralinear Ca^2+^ signals, observed when a CF-EPSP is preceded by PF activation, that are believed to trigger PF synaptic plasticity (Hartell, 2002; Jörntell and Hansel, 2006; Vogt and Canepari, 2010). In a recent report, we have demonstrated that isolated CF-EPSPs essentially trigger Ca^2+^ influx *via* T-type VGCCs when the initial V_m_ < −70 mV and Ca^2+^ influx *via* P/Q-type VGCCs when the initial V_m_ > −60 mV (Ait Ouares et al., 2019). In both cases these signals are spread throughout the dendritic arborisation at different extent. Here we demonstrate that bursts of PF-EPSPs are capable of locally amplifying these two Ca^2+^ signals at the sites of activated PF synapses, driving the CF “error” signal specifically to these inputs (Safo and Regehr, 2008). The biophysical mechanisms that permit these phenomena of amplification are summarised in Figure 12. As shown in Figure 6B, a burst of PF-EPSPs locally depolarises the dendritic area comprising the activated spines. Thus, if a CF-EPSP occurs during this depolarising phase, the Ca^2+^ influx associated with the CF-EPSP will be principally mediated by P/Q-type VGCCs, independently of the initial V_m_. The fast Ca^2+^ influx associated with the PF-mediated depolarisation can transiently saturate the endogenous Ca^2+^ buffer and therefore a supralinear free Ca^2+^ signal is produced in the dendritic region where V_m_ is depolarised. It must be pointed out that this type of supralinear Ca^2+^ signal depends on the number and arrangement of activated PFs that determine the spatial pattern of depolarisation, but it is not clearly established whether a scenario of “beams” of PF axons activating spines in close proximity occurs *in vivo* (Bower, 2002). During the short PF-mediated depolarisation, mGluR1s do not affect the Ca^2+^ transient associated with the CF-EPSP (Figure 1D). After the end of the PF train, V_m_ is repolarised and mGluR1 effects take action. In this time window, if the repolarised V_m_ is at the *hyp* state (< −70 mV) the Ca^2+^ influx associated with the CF-EPSP will be principally mediated by T-type VGCCs which are potentiated by mGluR1s (Figure 8). In addition, the slow mGluR1-mediated Ca^2+^ influx is also capable of saturating the endogenous buffer (Figure 9), a mechanism that further amplifies this supralinear free Ca^2+^ signal (Figure 9). Interestingly, both T channels potentiating and the slow mGluR1-mediated Ca^2+^ influx are triggered by the same mGluR1 pathway (Hildebrand et al., 2009) involving a protein tyrosine phosphatase (Canepari and Ogden, 2003), suggesting that the two factors contributing to this supralinear Ca^2+^ signal co-localise with mGluR1s at submicron scale. Eventually, if the repolarised V_m_ is > −65 mV, the Ca^2+^ influx associated with a CF-EPSP will be principally mediated by P/Q-type VGCCs. These channels can be also potentiated by mGluR1s through inhibition of A-type K^+^ channels (Otsu et al., 2014), but with two major differences with respect to the T channels boosting discussed above. First, this mechanism is highly variable since at these initial V_m_ range A-type K^+^ channels are already partially inactivated. Second, the potentiated P/Q channels do not co-localise with the site of activated PFs (Figure 10A-B). Thus, the only mechanism providing co-localisation in this case is saturation of the endogenous buffer produced by the slow Ca^2+^ influx that is independent of the initial V_m_.

### Role of the initial V_m_ and implication for synaptic plasticity

The evidence that mGluR1s locally potentiate a specific Ca^2+^ source (the T-type VGCC) and concomitantly amplify the same signal suggests that the occurrence of a *hyp* state episode is important for synaptic plasticity. Yet, *in vivo* electrode recordings have shown that, in PNs, V_rest_ is similar to that observed in brain slices (Kitamura and Häusser, 2011), indicating that the dendritic V_m_ is normally not at the *hyp* state. Also, spontaneous bursts of dendritic Ca^2+^ spikes are observed by recording Ca^2+^ and V_m_ signals simultaneously (Roome and Kuhn, 2018), again indicating that depolarising transients occur when the dendrite is relatively depolarised. These observations suggest that the *hyp* state might occur exclusively during rare but crucial episodes in which synaptic plasticity may occur. Episodes of *hyp* state can be due, in principle, to synaptic inhibition coactively occurring during activity patterns associated with cerebellar learning (Suvrathan and Raymond, 2018). Indeed, it was recently demonstrated that optogenetic activation of interneurons in the molecular layer strongly affects PF synaptic plasticity and motor learning, as well as CF-mediated signalling (Rowan et al., 2018; Gaffield et al., 2018). Another possibility is that *hyp* state episodes are intrinsically generated in PNs by a “priming” signal that hyperpolarises the cell, for instance by activating a K^+^ conductance. Interestingly, in a recent study, it was proposed that PF synaptic depression may physiologically occur when two CF events occur, the first one concomitant with a PF-EPSP burst and the second one after 100 ms (Bouvier et al., 2018). Notably, the concomitant activation of PF and CF inputs hyperpolarises the PN, as also visible in the examples with 60 ms delay reported here in Figure 3A and Figure 7A.

### Role of the mGluR1-activated non-selective cation conductance

An mGluR1-activated non-selective cation conductance, permeable to Ca^2+^, is the channel that mediates the slow mGluR1 Ca^2+^ influx (Canepari et al., 2001; Canepari et al., 2004). This channel has been identified as the C3-type transient receptor potential (TRPC3) since mutant mice lacking this channel all signals associated with the mGluR1-activated non-selective cation conductance are absent (Hartmann et al., 2008). This mouse, as well as another mouse model carrying a point mutation in the TRPC3 channel (Becker et al., 2009), exhibit severe changes in behavioural cerebellar functions. More recently, it was shown that mGluR1s can also activate GluD2 delta “orphan” glutamate receptors, raising the question of whether the mGluR1-activated non-selective cation conductance is actually composed by different channels (Ady et al., 2008). Notably, critical mutations of GluRD2 impair synaptic plasticity and motor learning and cerebellar behavioral functions (Hirano, 2006). Regardless of the identity of the channel underlying the slow mGluR1 Ca^2+^ influx, in this study we report a clear role for this conductance. During the time window of mGluR1 action, Ca^2+^ entering the cell after a PF-EPSP burst binds to mobile proteins forming the rich endogenous buffer of PNs (Bastianelli, 2003). Therefore, in this critical time window, the buffer capacity of PN dendrites, which is exceptionally high at rest (Fierro and Llano, 1996), can locally lower to a level that enables Ca^2+^ influx associated with a CF-EPSP to activate molecular pathways that would not be activated by a CF-EPSP alone. The transient saturation of the endogenous buffer has been already identified as crucial computational element that allows local integration of Ca^2+^ signals (Maeda et al., 1999). However, experimental evidence of endogenous buffer saturation was associated with dendritic depolarisation (Canepari and Vogt, 2008), co-localised with activated VGCCs which go beyond the spatial pattern of activated spines. In contrast, endogenous buffer saturation caused by the slow Ca^2+^ influx can be segregated to activated synapses. In general, the spatial pattern of activated spines (Schmidt et al., 2007) and their geometry (Schmidt and Eilers, 2009) are expected to be determinants of endogenous buffer saturation in the case of synaptic Ca^2+^ influx. Finally, endogenous buffer saturation caused by the slow mGluR1 Ca^2+^ influx is likely co-localising with the potentiated T-type VGCCs when the dendrite is at *hyp* state, since the two mGluR1-mediated mechanisms share the same molecular pathway (Hildebrand et al., 2009).

### Perspectives

This study was achieved by monitoring fluorescence simultaneously in large portions of PN dendrites with a spatial resolution of ∼10 µM. Thus, the dendritic sites of recording comprise several spines and the parent dendritic segment. A full understanding of the signals in terms of individual spines and dendritic bulks, which is not available in this study, should be achieved in the near future either by reducing the temporal resolution using rapid multisite confocal imaging (Filipis et al., 2018), or by reducing the number of recording spots using 2-photon random access microscopy (Otsu et al., 2008). A second question that should be explored in detail is the relation between the initial V_m_ state and the induction of PF synaptic plasticity. Numerous studies have investigated other parameters as crucial determinants of PF synaptic plasticity, including timing, size of Ca^2+^ signals, spatial arrangements of synaptic inputs and others (Vogt and Canepari, 2010). In this study we show that a scenario, where a specific Ca^2+^ source (the T-type VGCC) is locally boosted and amplified by mGluR1s, occurs only when the initial initial V_m_ is hyperpolarised, a phenomenon that can translate into an important information processing rule associated with cerebellar function.

## Acknowledgements

This work was supported by the *Agence Nationale de la Recherche* through grants ANR-14-CE17-0006 - WaveFrontImag; Labex *Ion Channels Science and Therapeutics:* program number ANR-11-LABX-0015 and National Infrastructure France Life Imaging “Noeud Grenoblois”); and by the *Federation pour la recherché sur le Cerveau* (FRC – Grant *Espoir en tête*, Rotary France). We thank Dr. Boris Barbour for useful discussions during the project and for reading the manuscript before submission.

